# Correlated structure of neuronal firing in macaque visual cortex limits information for binocular depth discrimination

**DOI:** 10.1101/844514

**Authors:** Jackson E. T. Smith, Andrew J. Parker

**Affiliations:** Department of Physiology, Anatomy and Genetics, University of Oxford, Oxford, Oxfordshire, OX13PT, United Kingdom

**Keywords:** V1, V4, Cortex, Vision, Rhesus, Noise correlation, Stereo depth, Encoding, Behaviour, Utah array

## Abstract

Variability in cortical neural activity potentially limits sensory discriminations. Theoretical work shows that information required to discriminate two similar stimuli is limited by the correlation structure of cortical variability. We investigated these information-limiting correlations by recording simultaneously from visual cortical areas V1 and V4 in macaque monkeys, performing a binocular, stereo-depth discrimination task. Within both areas, noise correlations on a rapid temporal scale (20-30ms) were stronger for neuron-pairs with similar selectivity for binocular depth, meaning that these correlations potentially limit information for making the discrimination. Between-area correlations (V1 to V4) were different, being weaker for neuron pairs with similar tuning, and having a slower temporal scale (100+ms). Fluctuations in these information-limiting correlations just prior to the detection event were associated with changes in behavioural accuracy. Although these correlations limit the recovery of information about sensory targets, their impact may be curtailed by integrative processing of signals across multiple brain areas.

## INTRODUCTION

Processing in the sensory areas of cerebral cortex is thought to limit the detection and discrimination of weak sensory signals (A. J. Parker & Newsome, 1998). Initial work examined how the changes in the firing of action potentials induced by near-threshold changes in the sensory stimulus could be detected against the variability of the base-level of cortical activity (Barlow, Kaushal, Hawken, & Parker, 1987; Newsome, Britten, & Movshon, 1989; A. J. Parker & Hawken, 1985; Tolhurst, Movshon, & Dean, 1983). This work elucidated how detection and discrimination could be related to the statistics of the firing of single neurons.

These initial studies indicated that it was not unusual for the detection performance of single neurons to match the behavioural performance of the entire visual system. However, since the firing of all neurons shows some variability, combination of the firing from a pool of neurons to form a multi-neuron sensory signal ought to improve the detection rate for weak stimuli. The puzzle is why the nervous system apparently cannot provide a better sensitivity for the organism by making a statistical combination of the signals from multiple neurons.

One answer from early work on the somatosensory periphery is that the correlation of residual noise within a pool of sensory neurons may also be an important factor (Johnson, Darian-Smith, & LaMotte, 1973). In the absence of correlations, additive pooling of neuronal signals results in a steady improvement of sensory thresholds with increasing pool size. However, the presence of correlations limits the potential gain in performance from increases in the numbers of neurons. As more and more neurons are added, the portion of noise that is uncorrelated does indeed get averaged away, but the portion of noise that is correlated, and therefore shared by every neuron, cannot be reduced this way and comes to dominate the limit on performance (A. J. Parker & Newsome, 1998; Shadlen, Britten, Newsome, & Movshon, 1996).

### Correlation structure in the pool of sensory neurons

Recent work expands that insight, showing how the nature of the pooling computation and the structure of correlations are both critical in setting limits on the encoding and decoding of information within sensory cortex (Abbott & Dayan, 1999; Averbeck, Latham, & Pouget, 2006; Kanitscheider, Coen-Cagli, Kohn, & Pouget, 2015b; Moreno-Bote et al., 2014; Pouget, Dayan, & Zemel, 2003). The correlation structure of a pool of neurons in relation to sensory discriminations is captured by making all pair-wise comparisons between members of the pool, across a number of repeated presentations of the sensory stimuli. Critically for our study, this theoretical work demonstrates that the signals from a neuron-pair limit the discrimination between two stimuli in two ways (Nogueira et al., 2020). The first is fairly straightforward, which is that the change in the stimulus should alter the mean level of firing from both neurons; bigger differences in the mean response should improve the discrimination ability. The second concerns the variability of neuronal firing and its correlation structure across the population. The theoretical work demonstrates that noise or variability is the most detrimental when its correlated structure causes the conjoint firing of the neuron-pair to mimic a response to some change in the stimulus that did not, in fact, occur. Such correlations have been termed “information-limiting correlations” (Kohn, Coen-Cagli, Kanitscheider, & Pouget, 2016; Moreno-Bote et al., 2014).

Assessment of the correlation structure ideally requires the simultaneous recording of multiple pairs of neurons from within the sensory cortex during the performance of a behavioural task with repeated stimulus presentations (Kanitscheider et al., 2015b; Moreno-Bote et al., 2014). In our work, we addressed this by training monkeys to discriminate binocular stereoscopic depth, while we tested a range of neuronal encoding by using chronically implanted Utah arrays to record multiple neurons simultaneously in two cortical sites: the primary visual cortex (V1) and the extrastriate area V4. Although the fact that we studied binocular vision is not especially critical for the questions that we investigate here, the processing of stereoscopic information proves to be a good model system for isolating the contribution of cortical processing, as stereo involves the direct combination of information from the two eyes at a cortical level.

### Neuronal recording during sensory task performance

Our task was designed so that we could thoroughly test all simultaneously recorded neurons over a range of stereoscopic depths, to build up a complete tuning curve for each neuron. Many studies have employed tasks that involve a binary decision, potentially limiting which cortical neurons are relevant to the task, with the result that many of the recorded neurons may never be tested under the optimal (or near-optimal) conditions for assessing their encoding of information. Measurement over the whole range of sensory encoding also increases the yield of relevant neuronal responses in the case of using fixed recording electrodes, as with the Utah array.

The paradigm used here allowed us to estimate the extent to which the two neurons respond in the same way to a range of stimulus values, which is termed the signal correlation (Bair, Zohary, & Newsome, 2001) for the neuron pair. As our task involved the discrimination between two closely similar binocular depths, we also assessed the differential response of the sensory neurons by taking the slope of the tuning curve at each point. By making repeated measures of the response to each binocular depth, we could also estimate the noise correlation of all recorded neuron-pairs at each tested value of binocular depth, and apply recently-developed methods to search for information-limiting correlations (Kanitscheider, Coen-Cagli, Kohn, & Pouget, 2015a). We couple this approach with a time-dependent measure of correlations (Bair et al., 2001) to estimate the temporal scale of information encoding.

In earlier comparisons of the correlation structure and sensory tuning of cortical neurons, the data were either from anaesthetized animals (M. A. Smith & Kohn, 2008) or from awake animals that fixated without performing any other task (M. A. Smith & Sommer, 2013). In other cases, the behavioural measures were taken on one task (judgment of heading direction) but the measures of correlation structure were taken during the performance of a different task (visual fixation) (Pitkow, Liu, Angelaki, DeAngelis, & Pouget, 2015). In an alternative scenario, the measurements of signal correlation were obtained whilst the animal was passively fixating rather than actively discriminating the stimuli (Bondy, Haefner, & Cumming, 2018).

An important advance made with the recordings undertaken here is that the macaque monkeys actively performed a behavioural task, making a stereoscopic depth judgment on the presented stimuli. We obtained simultaneous neuronal and behavioural data across the complete tuning curve of the neurons. A number of recent studies have demonstrated that the correlation structure of neuronal firing in sensory areas is influenced by the manipulation of cognitive factors such as attention or perceptual learning (Bondy et al., 2018; Ni, Ruff, Alberts, Symmonds, & Cohen, 2018; D. A. Ruff & Cohen, 2014; Douglas A. Ruff & Cohen, 2016; Verhoef & Maunsell, 2017). We aimed to maintain these factors as constant as possible to focus on the discrimination performance of the neuronal population, by ensuring that the design of the experimental task arranged for spatial and featural attention to be consistently directed towards the depth signalled by the visual stimuli.

Finally, the relationship between perceptual decisions and the correlation structure of neuronal firing has been a topic of intensive research (Krause & Ghose, 2018; Krug, Curnow, & Parker, 2016; Pitkow et al., 2015) ever since it was suggested that the two may be linked (Shadlen et al., 1996). To assess this link, it is critical to measure the correlation structure in the most relevant neuronal population as the animal is performing a sensory discrimination task. In this paper, we are primarily concerned with the state of cortical firing as the animal waits attentively for the arrival of a change in binocular depth that must be detected and reported. Our results show for the first time that the state of putative differential correlations within the neuronal population has a direct association with the accuracy of the upcoming behavioural decision.

### Correlations within and between cortical areas

A further open question is whether the noise correlation structure limits the passage of signals through each of the stages of cortical processing (Zylberberg, Pouget, Latham, & Shea-Brown, 2017). We tackle this issue by examining the correlation structure and information encoding of neuronal activity within and between cortical areas V1 and V4. Specifically, we test how information may pass between cortical areas, with respect to neuronal correlations during task performance. We examine the time-course of information-bearing signals and assess them in relation to information-limiting correlations and behavioural choice. Our results suggest a simple model, which demonstrates that the potentially deleterious effects of information-limiting correlations do not build up as sensory signals pass from one cortical area to another. We suggest that this is an important and previously unrecognized principle of neural encoding across multiple areas of the cerebral neocortex.

## RESULTS

We trained two Rhesus macaques (*Macaca mulatta*) to observe 4 patches of dynamic, random dot stereograms (RDS) and perform an odd-one-out task (Figure 1), in which they detected which one of the four patches displayed a change in binocular depth. Each RDS was configured as a central circular patch, whose binocular depth varied from trial to trial, and a surround region of dots located in the fixation plane of the monkeys’ eyes. In the ‘present RDS’ phase (Fig.1Aiii), all 4 stimuli had the same depth for the central patch, which was chosen from one out of a set of 13 different values. The ‘popout’ phase (iv) presented the change in depth, which was applied to the centre of one of the 4 RDS stimuli. That patch became the target for a saccade to indicate a correct choice and earn a reward; a saccade to any of the other three RDS was incorrect and earned no reward.

**Figure 1.**
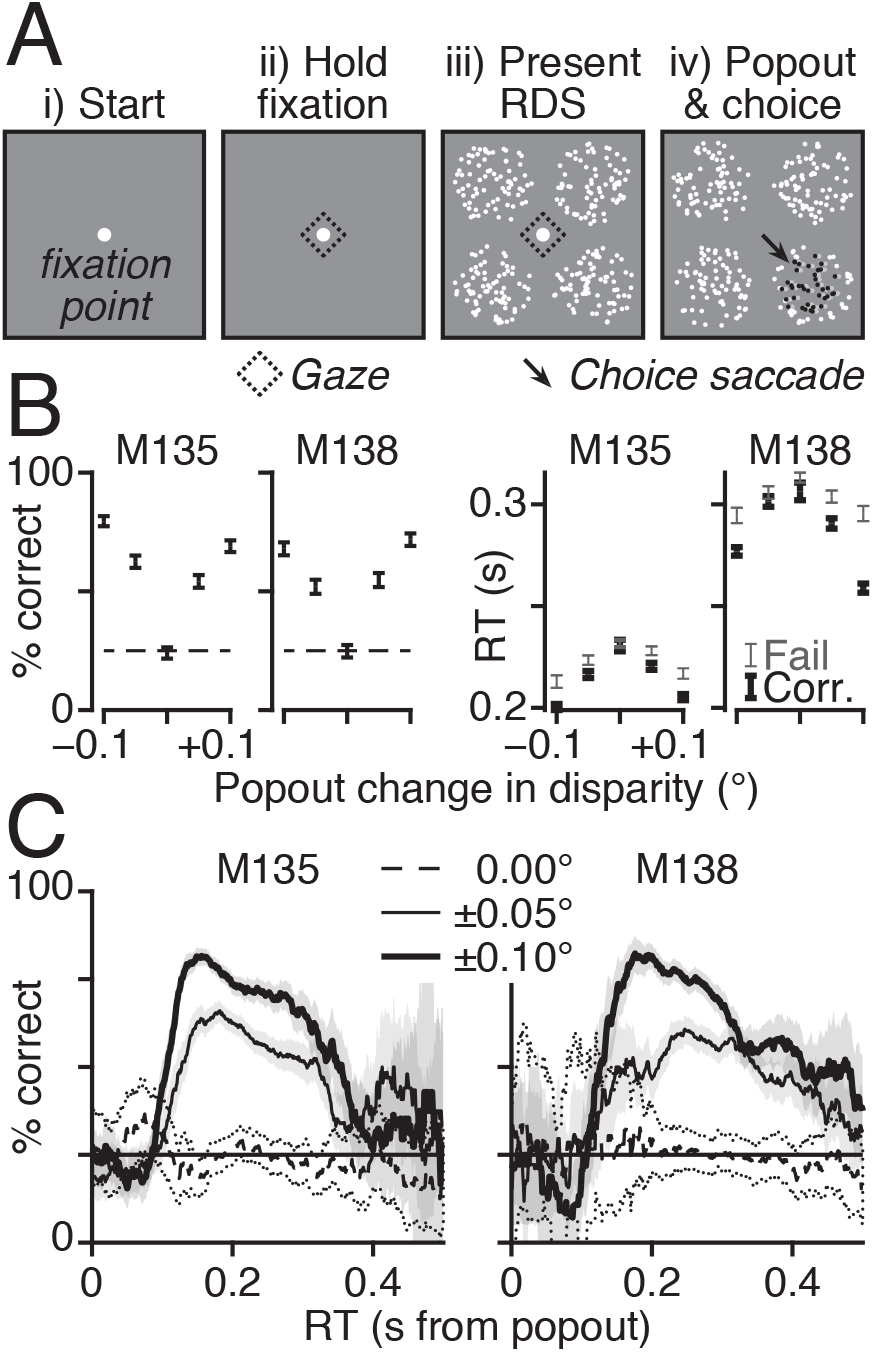
Task and performance. A) (i) Trials started with flashing fixation point (FP, white circle). (ii) Once fixated (dashed diamond), FP flash stops. (iii) 500ms later, four random dot stereograms (RDS, white dots) with identical properties appeared. (iv) After fixed duration (1s M135, 2s M138), central dots of one RDS change disparity (grey/black dots) and FP disappears. Animals made saccade onto the oddball target RDS (arrow) for reward. B) Percent correct trials (chance=25%, dashed line) and reaction time (RT) over popout disparity. Using trials with 0.117 ≤ RT ≤ 0.344s for M135 (N = 6670 of 8279), or 0.143 ≤ RT ≤ 0.463s for M138 (N = 5478 of 6073). RT shown separately for correct (thick) and incorrect choices *i.e*. failed trials (thin). Error bars 95% binomial confidence intervals (CI) or standard error of the mean (SEM). C) Percent correct across RT using 50ms moving window; with 95% bootstrap CI (2000 samples, 2.5^th^ & 97.5^th^ percentiles). Data grouped by absolute popout disparity (see legend).

The monkeys’ behaviour indicated attention to the RDS stimuli in the ‘Present RDS’ phase, as they were better at reporting the location of the disparity step change as it grew in magnitude (Fig.1B, % correct), while their reaction times (RT) decreased. On catch trials with no change in the RDS, the animals had to guess, so their accuracy was at chance levels (Fig.1B, dotted line, left panels) and their RTs were longest (Fig.1B, right panels). Fig.1C shows the relationship between percent correct as a function of RT using a 50ms moving window, grouped by magnitude of the popout disparity change. Correct decisions are broadly within the window of 100-400ms after pop-out.

Neural activity in the dorsal, gyral portions of V1 and V4 was recorded, using two 64 channel (8×8; 0.4mm spacing) Utah array electrodes (one array per area) implanted chronically in the left hemisphere. The electrode placement meant that the recorded neurons had receptive fields (RF) that were stimulated by the lower-right RDS (Figure 2A&S1). An automatic spike sorting method was used to identify action potentials followed by manual checking (see Methods for details.) As with other work with Utah arrays, analyses were based on both single units and multiunit clusters and the term ‘unit’ refers interchangeably to either.

**Figure 2.**
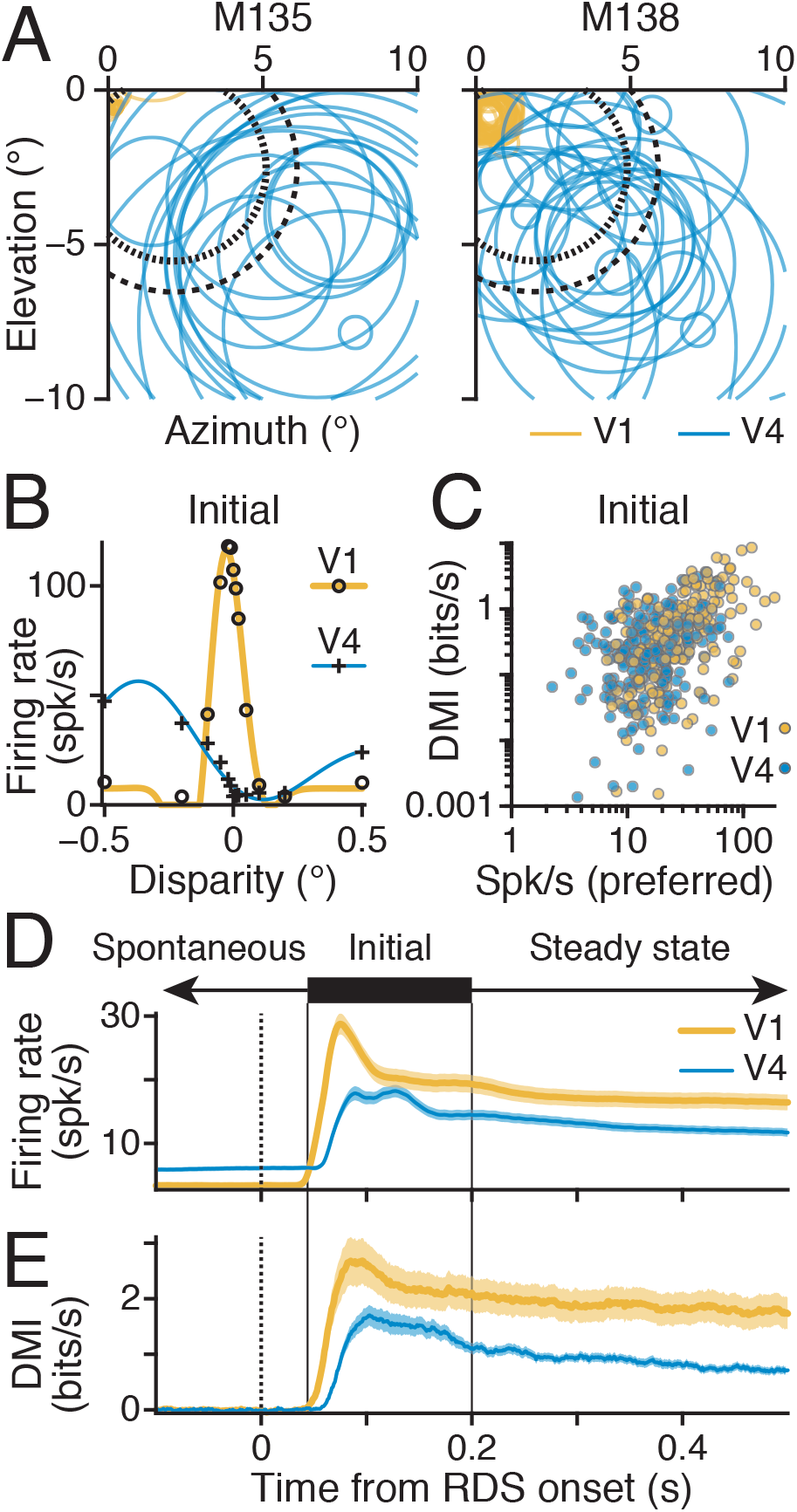
V1 and V4 units were selective for binocular disparity. A) Receptive field (RF) location and size for each V1 (gold) and V4 (blue) neurons; circle radius = 25% standard deviation of 2D Gaussians fit to RF location maps. RDS centre (short dash) and surround (long dash) shown. B) Example disparity tuning curves from V1 (circles, yellow) and V4 (crosses, blue); symbols show average firing rates; lines show best-fitting Gabors: SEM below symbol size. C) Disparity mutual information (DMI) for V1 (gold, N = 195) and V4 (blue, N = 213) units compared with average firing rate to preferred disparities. D) Average firing rate (mean line, SEM ribbon) across all V1 (gold) and V4 (blue) units, over trial sequence: analysis windows cover spontaneous responses (left arrow), Initial phasic response (solid bar and lines), later Steady state responses (right arrow); dashed line marks the onset of RDS. E) As in C but for DMI.

Variations between neuronal RF with respect to spatial location and preference for binocular depth, dot size, dot contrast and other dimensions of the RDS stimulus mean it is impossible to simultaneously achieve optimal stimulation of every unit recorded by the arrays. We therefore focused on varying these parameters to ensure that there was significant co-activation of the V1 and V4 units (Fig.2D&E, S2D&E). To validate the co-activations of neuron pairs, we applied entry criteria, such that each unit in the pair had to show statistically significant tuning to disparity (one-way ANOVA at the 1% level) and fire at least 5 spikes per second to its preferred disparity. For all the recordings reported here, the animals were performing the stereo depth task, which requires allocation of attention to the four RDS patches.

We made separate measurements of RF location and stimulus preferences in preliminary experiments in each monkey. These measurements demonstrated that there was scatter in the positions of the RF centres of the units, both within and between cortical areas (Fig.2A&S1). We later address how this separation of RF centres may have influenced our results. For now, we confine ourselves to noting that the combination of electrode placements and adjustment of the configuration of the RDS stimulus resulted in common activation of pairs of V1 and V4 units by the centre portion of the same RDS. We focus on the activations of units from the ‘Hold fixation’ and ‘Present RDS’ phases of the trial (Fig.1Aii&iii, 2D&E), before the arrival of the discrimination stimulus itself.

### V1 and V4 units were selective for binocular disparity

The V1 and V4 neural signals that we recorded are able to support task performance. Disparity tuning curves (Fig.2B) are shown for an example V1 (circles, gold line) and V4 (plus sign, blue line) unit. The solid lines are the best-fitting Gabor functions (equation 21)(Prince, Pointon, Cumming, & Parker, 2002), which describe the response profiles as a function of disparity. To quantify a unit’s contribution to the estimation of stimulus disparity, we computed how much information the firing of the neuron could contribute to the identification of the currently displayed disparity of the RDS, called the disparity mutual information (DMI, equation 8; Fig.2C&E, S2A-C&E). This measure correlated well with a previously-established measure of selectivity (Fig.S2A&B) – the disparity discrimination index (DDI; Prince et al., 2002) – for the V1 (Spearman correlation, Initial, *r* = 0.976, *p* = 9.559e–130) and V4 (*r* = 0.953, *p* = 4.705e–111) neurons.

The neurons in V1 and V4 that responded most to the onset of the stimulus were also the most selective for disparity, as the peak firing rate and DMI (in the 50 to 200ms window) were well correlated (Fig.2C&S2C: V1 in gold, Spearman *r* = 0.609, *p* → 0; V4 blue, *r* = 0.248, *p* = 2.663e–4). Low-firing neurons were relatively poor at encoding disparity, compared to highly active neurons.

We searched for the time point at which the units became selective for disparity by convolving the spike trains with a causal exponential kernel (20ms time constant) to obtain the firing rate and DMI time series for each unit. The average firing rates (Fig.2D&S2D; V1, gold; V4, blue) had a transient burst of activity in response to the onset (black dotted) of the RDS, which then settled down to a steady rate. As expected, V1 began responding to the RDS before V4 and V1 neurons showed selectivity for the binocular disparity slightly earlier than V4 neurons.

Firing rates in both areas were selective for binocular disparity almost as soon as the units began responding to the RDS, with the average DMI showing a similar time course to the changes in the firing rate (Fig.2E&S2E). We verified the link in timing between the DMI and the peak firing for each unit by finding the Spearman correlation of firing rate and DMI, matched by time bin, in an 800ms window starting 50ms after the RDS onset *i.e*. we took the zero-lag cross-correlation of the firing rate and DMI time series. The result was significantly positive for V1 (mean *r* = 0.192, right-tailed t-test *p* = 3.339e–22) and V4 (*r* = 0.245, *p* = 7.819e–31), showing that individual units were more selective for disparity at moments in the trial when their firing rate was higher.

We partitioned each trial into three consecutive windows for further analysis (Fig.2D&E). The ‘Spontaneous’ window (leftward arrow) captured neural responses during the ‘Hold fixation’ phase (Fig.1Aii) of the trial, in which the monkeys held their gaze on the fixation target prior to the arrival of RDS signals in V1. The response to the onset of the RDS is captured by the ‘Initial’ window (thick bar and black solid lines, marking 50-200ms after stimulus onset), which shows a transient burst of firing and rise in the DMI. The responses to the RDS for the remainder of the ‘Present RDS’ phase of the trial are captured by the ‘Steady state’ window (rightward arrow, Fig.1Aiii). Within-unit comparisons showed that disparity selectivity (DMI) was stronger in the ‘Initial’ than in the ‘Steady state’ window, for V1 (mean paired difference = −0.577 bits/s, left-tailed t-test *p* = 1.294e–12) and V4 (−0.220 bits/s, *p* = 6.827e–11); in part, these differences reflect the greater firing rates in the Initial window.

### Information-limiting correlations

If the sensory discrimination task performed by the monkeys is supported by a linear computation across pools of active neurons, then a special class of noise correlation – differential correlation (Moreno-Bote et al., 2014) – is potentially a fundamental limit on the information that is encoded by the sensory pool. Figure 3A–C show how differential correlations could limit information that is retrievable from a pair of neurons, whose activities were simultaneously recorded in V4 during task performance. The pair had similar tuning for binocular disparity (*i.e*. 0 < *r_signal_*, Fig.3A). Both units had a common negative slope in their tuning curves around 0.1° (black dashed), where they were sensitive to the same small changes in disparity. Adding the signals from these two neurons potentially improves sensitivity but this depends on the noise distributions.

**Figure 3.**
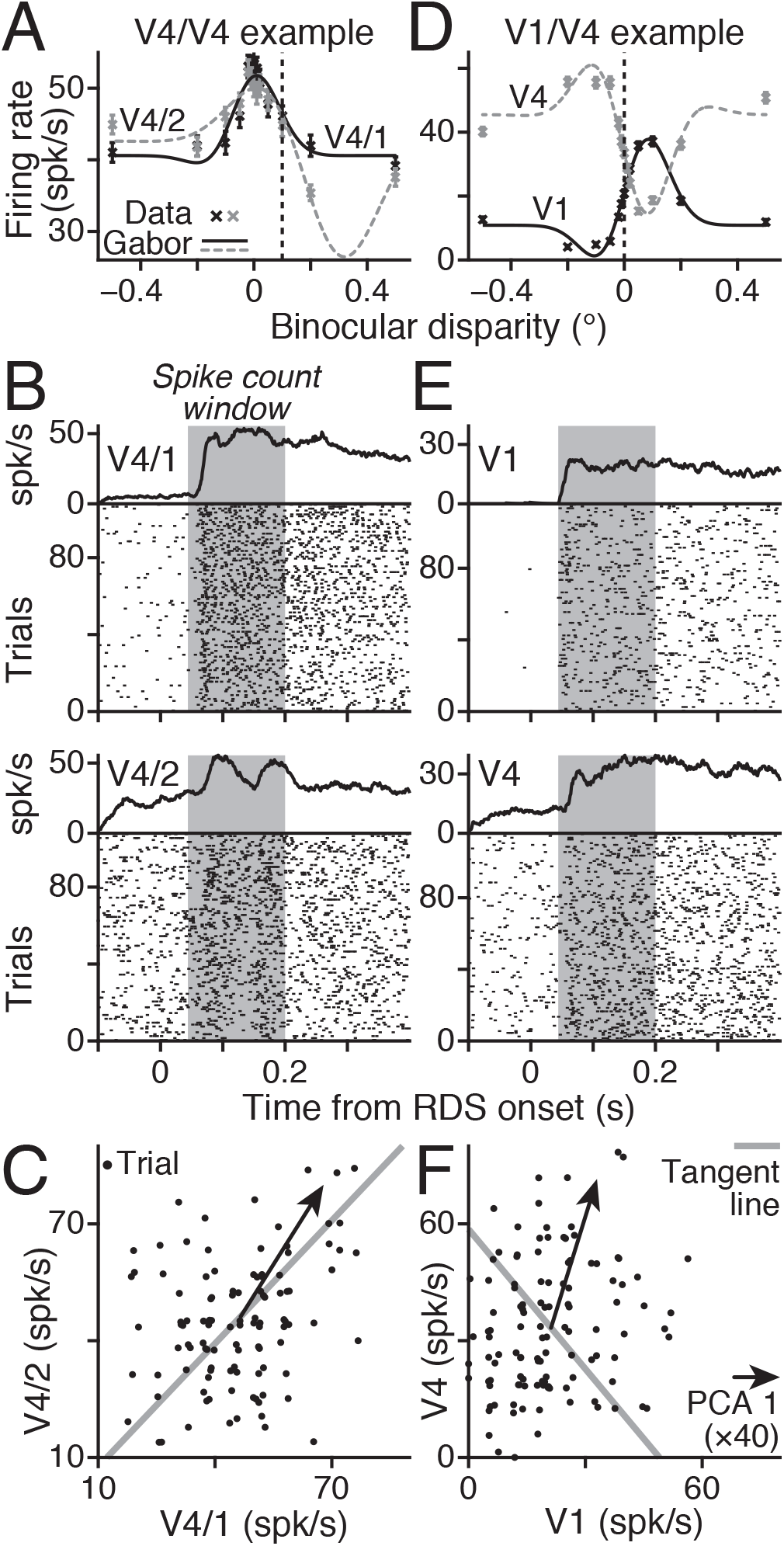
Alignments of noise correlations and disparity tuning. A) Disparity tuning curves of pair V4/1 (black) & V4/2 (grey; M138.336.94/147), signal correlation (*r_signal_* = +0.851); crosses and SEM show average firing, lines show best-fitting Gabors (V4/1 *r*^2^ = 89%, V4/2 *r*^2^ = 93%). Vertical dashed line at +0.1° shows trials (N = 108) examined in panels B&C. B) Raster plots and average firing rates for V4/1 (top) and V4/2 (bottom); RDS onset at t=0; for 108 trials at baseline disparity +0.1°; shading shows time window for firing rates in C. C) Dots show joint firing rates; slight jitter applied for visualization; noise correlation, *r_SC_* = 0.241, *p* = 0.012. Tangent line (grey) shows joint tuning curves at +0.1°; arrow shows first PCA component (×40, for visibility) of joint responses. D – F) the same as A – C but for an example V1/V4 pair (M138.336.57/87; V1 black in D, *r*^2^ = 98%; V4 grey in D, *r*^2^ = 94%) with negative signal correlation (*r_signal_* = −0.982). Trials with baseline disparity of 0° (D, black dashed, N = 116) yield significant positive noise correlation (F, *r_SC_* = 0.203, *p* = 0.029).

Joint responses to multiple presentations of the 0.1° disparity stimulus were obtained by counting spikes in the Initial window (Fig.3B, grey). At 0.1°, the pair had a positive noise correlation (Fig.3C). Critically, the principal component (arrow) of the covariance was nearly parallel to the tangent line (grey) of the tuning curves at 0.1°. To see the consequence of this, consider a linear discriminator that classifies the joint responses as one of two disparities on either side of 0.1°. If the decision boundary is perpendicular to the tangent line (grey) then noise correlation (arrow) will make the joint responses jump across the decision boundary, causing classification errors (Averbeck et al., 2006). Put simply, correlated noise would mimic true changes in the stimulus, limiting the sensitivity of this pair.

This outcome contrasts with the responses of a second pair (Fig.3D–F), recorded simultaneously from V1 and V4. As before, both units had steep, overlapping slopes in their tuning curves (Fig.3D). Unlike the V4/V4 pair, the V1 and V4 units were selective for opposite disparities (*r_signal_* < 0). Spike counts from both units in response to a disparity of 0° (Fig.3D, black dashed) were obtained as before (Fig.3E), and the joint responses were examined (Fig.3F). Although this pair has a negative signal correlation, they have a positive noise correlation (*r_SC_* = 0.203; p = 0.029): the noise correlation does not align to the tangent (noise correlation first principal component PCA1 at 72.8° [95%CI: 58.5°, 85.3°]). In this case, correlated noise would be less likely to leak across the decision boundary. Further examples of neuron-pairs with opposite signs of relationship between signal and noise correlation are shown in Supplementary Figure S3. For those examples, one pair also had opposite selectivity and positive correlation, but two others had the inverse pattern, with similar disparity selectivity and negative noise correlations.

### r_CCG_ measures noise correlation at different time scales

Noise correlations (*r_SC_*, equation 10&11) are captured experimentally by setting up a window of time, in which to count the number of spikes in response to repeated presentations of the same stimulus. In terms of timing, this encompasses the synchronous firing of spikes with millisecond precision up to long time-scale, co-fluctuations in spike numbers (Marlene R. Cohen & Kohn, 2011; M. A. Smith & Kohn, 2008; M. A. Smith & Sommer, 2013). Measurement of spike count correlation confounds these slow and rapid correlations, depending on the size of the counting window that is used. Another, time-based measure – *r_CCG_* (equation 17) – is used to reveal noise correlations at different temporal scales (Bair et al., 2001). This measure, *r_CCG_*, is the time integral of the spike-train cross-correlation and reveals the timing relationships between correlated spikes, as well as the strength of the correlation. The *r_CCG_* measure converges on *r_SC_* as the width (*τ*) of the integration window reaches that of the counting window used to calculate *r_SC_*. We asked whether information-limiting correlations occupy a limited set of temporal scales by using *r_CCG_* to examine noise correlations within cortical areas V1 and V4 and between those two cortical areas (Figures 4 & S4).

**Figure 4.**
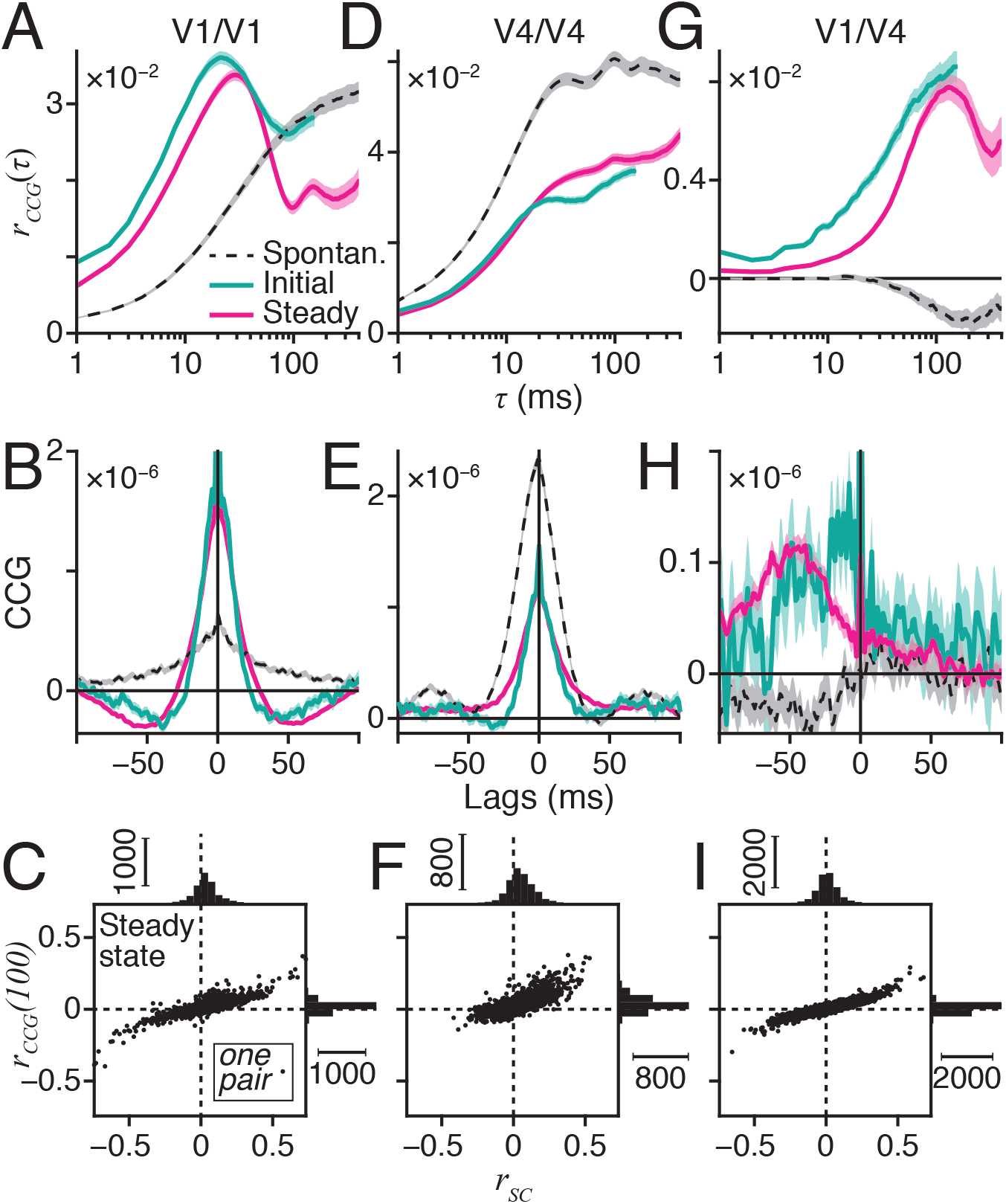
r_CCG_ measures noise correlation at different time scales. A) Average *r_CCG_* as function of temporal scale *τ* for all V1/V1 pairs’ (N = 2419) spontaneous (dashed black), Initial (teal), and Steady state (magenta) responses (see Fig.2D); ribbons show SEM. B) As A, but has average V1/V1 cross-correlograms (CCG). C) Scatter plot of Steady state *r_CCG_* at *τ* =100ms versus spike-count correlation (*r_SC_*). Marginal distributions show each metric separately. D – F) the same as A – C but for V4/V4 pairs (N = 2226). G – I) the same as A – C but for V1/V4 pairs (N = 4598).

Within V1 (Fig.4A), the time scale of noise correlations changed as the trial progressed. Before the RDS appeared, correlations occupied a slow time-scale, being strongest on the order of hundreds of ms (dashed). The time window for correlations became tighter after RDS onset, peaking at *τ* = 20ms; *r_CCG_*(τ=20ms) was greater in the Initial window (teal) than the Spontaneous window (mean paired difference = 2.141e–2, paired t-test *p* = 1.592e–124). Even in steady-state responses to the RDS (magenta), the slower correlations were weaker, with a drop in *r_CCG_* (τ=100ms) from the Spontaneous window to the Steady state (−1.092e–2, *p* = 1.005e–21).

These changes in *r_CCG_* over time scale are reflected in the shapes of the V1-V1 cross-correlograms (CCG, equation 19; Fig.4B). In the period of Spontaneous activity, the CCG was positive and had a broad peak, coinciding with the steady increase of Spontaneous *r_CCG_* as *τ* increased. In contrast, the Initial and Steady state CCGs had narrow positive peaks flanked by negative side lobes, showing that the majority of correlated spikes occurred within tens of ms of each other. This matched the peaks in Initial and Steady state *r_CCG_* at *τ* = 20ms.

Unlike V1, correlations within V4 (Figs.4D & S4A) began saturating at *τ* = 20ms in all three analysis windows. V4 correlations were also stronger in the Spontaneous window, with *r_CCG_*(τ=20ms) dropping in the Initial window (−2.188e–2, *p* = 9.053e–105) and then changing little from the Initial to Steady state windows (9.464e–4, *p* = 0.041). These differences appeared in the V4 CCGs (Figs.4E & S4B) in which the Initial and Steady state CCGs were mainly attenuated versions of the Spontaneous CCG.

In marked contrast, the correlations between V1 and V4 (Fig.4G) saturated at *τ* = 100ms. Although these between-area correlations are weaker, it is important to understand that they were significantly positive during the stimulus presentation (mean *r_CCG_*(100ms); Initial = 8.075e–3, right-tailed t-test *p* = 1.512e–43; Steady state = 7.398e–3, *p* = 4.088e–61; Spontaneous = −1.286e–3, *p* = 1). There was no measurable change in the V1/V4 correlations from the Initial to the Steady state window (mean difference = −6.767e–4, paired t-test *p* = 0.159). The V1/V4 CCGs (Fig.4H) reveal a consistent delay, whereby V1 spikes tended to occur up to 50ms earlier than the associated V4 spikes.

Bair and colleagues (Bair et al., 2001) established an important property of *r_CCG_*, which is that it provides a lower-variance estimate of short-term correlations, compared to *r_SC_*. The parameter *τ* can be adjusted so that *r_CCG_*(*τ*) integrates only the central peak of the CCG and discards the longer time-course components. These longer time-course components are captured by *r_SC_* but often add variance to that estimate. This principle is illustrated for the “Steady state” condition by a direct comparison of *r_CCG_*(100ms) and *r_SC_* in V1 (Fig.4C), V4 (Fig.4F&S4C), and between areas (Fig.4I). There was a strong, positive Spearman correlation of *r_CCG_* and *r_SC_* in all cases (V1 *r* = 0.750, V4 *r* = 0.772, V1/V4 *r* = 0.807, all *p* → 0), showing that they measured a common source of correlation. However, the marginal distributions plotted at the top and right-hand sides show that the variation (standard deviation) of *r_SC_* (*σ_SC_*) was greater than that of *r_CCG_*(100ms) (*σ_CCG_*) for all three cases (V1, V4, V1/V4, see Table 1). Hence, the Pitman’s *r_SD_* test for comparing variances of correlated variables (Lee, 1992) shows that *r_CCG_* is a more accurate measure of short-term correlations than *r_sc_*.

**Table 1:** Comparison of scatter in estimates of r_sc_ and r_ccg_(100ms)

| Cortical Area | $\sigma_{SC}$ | $\sigma_{CCG} (\tau=100\text{ms})$ | Pitman’s $r_{SD}$ |
| --- | --- | --- | --- |
| V1 | 0.135 | 0.044 | 0.830 |
| V4 | 0.112 | 0.056 | 0.766 |
| V1/V4 | 0.112 | 0.030 | 0.886 |

Each $r_{SD}$ has a p-value approaching 0

### Noise correlations varied with disparity tuning, temporal scale, and time

We asked initially whether the noise correlations of these neurons had any consistent relationship to disparity selectivity (Figure 5 & S5). We plotted *r_CCG_* as a function of the signal correlation between pairs (*r_signal_*, equation 9), grouped by temporal scale (*τ*, Fig.5A,C,E&S5A), or as a function of *τ*, grouped by *r_signal_* (Fig.5B,D,F&S5B).

**Figure 5.**
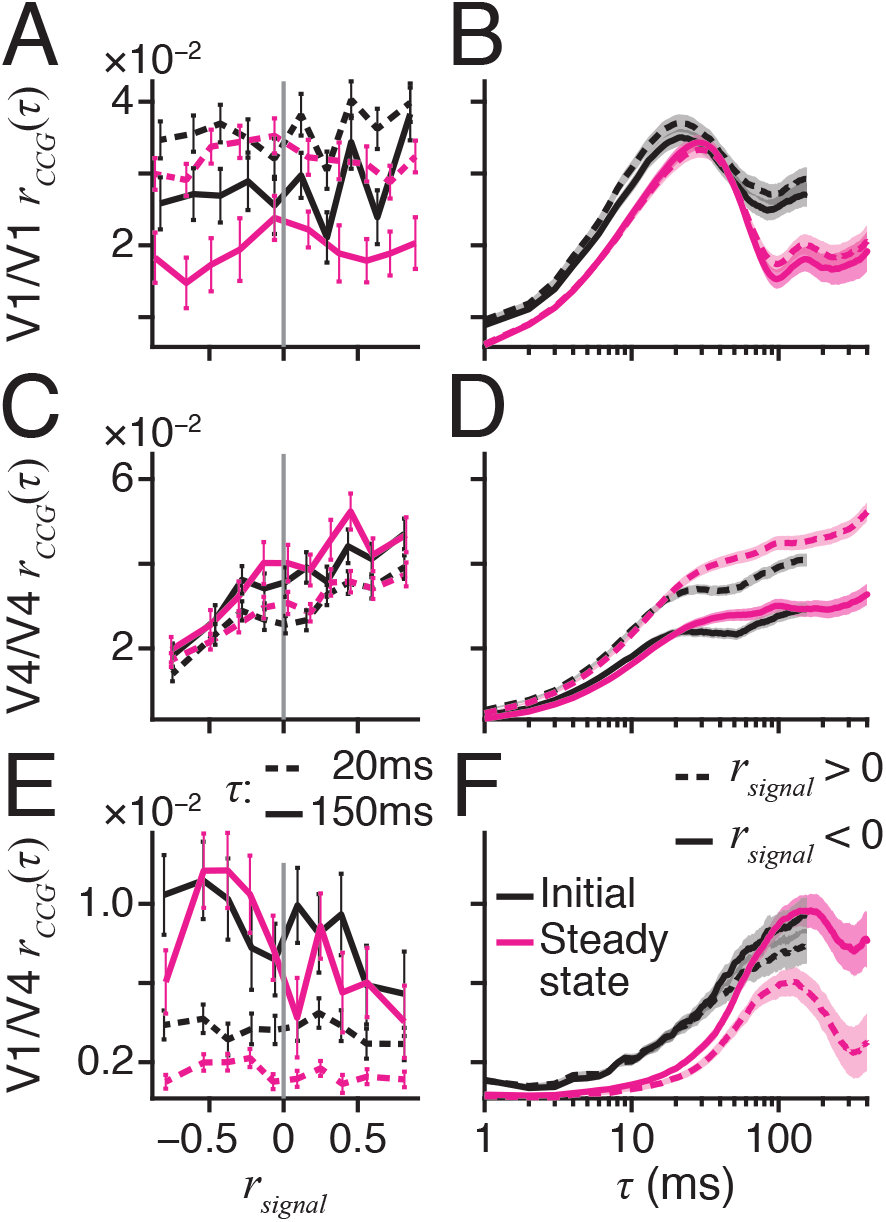
Noise correlations varied with disparity tuning, time scale, and time. A) Noise correlation (*r_CCG_*) for V1/V1 as function of signal correlation (*r_signal_*), grouped by time scale (*τ*) of *r_CCG_* (20ms, dashed; 150ms, solid) and analysis window (Initial, black; Steady state, magenta); each bin has 10% of the data; error bars are SEM. B) Average V1/V1 *r_CCG_* over time scale *τ*, grouped by signal correlation (positive *r_signal_*, dashed; negative *r_signal_*, solid) and analysis window, as A; ribbon shows SEM. C&D) same as A&B but for V4/V4 pairs. E&F) same as A&B but for V1/V4 pairs.

For pairs in V4, there was a clear positive relationship of *r_CCG_* and *r_signal_* (Fig.5C&S5A) at both short (*τ* = 20ms, dashed) and long (*τ* = 150ms, solid) time scales in the Initial (black) and Steady state (magenta) windows (see Table 2). This coincided with an increase in *r_CCG_* across a range of values of *τ* for pairs with similar disparity preferences (Fig.5D&S5B, *r_signal_* > 0, dashed), compared to those with opposite preferences (*r_signal_* < 0, solid). For pairs in V1, a similar but weaker positive relationship was observed (Fig.5A&B) that was significant in the Initial window and Steady state window for long time scales (*τ* = 150ms) but not short (*τ* = 20ms) (see Table 2). In summary, neurons in the same cortical area that have similar preferences for the binocular depth of the stimulus tend also to show stronger, more positive noise correlations, suggestive of a local network of neurons that process binocular depth.

**Table 2:** relationship between r_signal_ and r_ccg_, accompanies Figure 5

| Cortical areas | $\tau$ (ms) | Initial | | Steady state | |
| --- | --- | --- | --- | --- | --- |
| | | Spearman $r$ | P-value | Spearman $r$ | P-value |
| V1/V1 | 20 | 0.042 | 0.038 * | 0.030 | 0.144 ns |
|  | 150 | 0.050 | 0.013 * | 0.054 | 0.009 ** |
| V4/V4 | 20 | 0.172 | 4.133e-16 *** | 0.170 | 5.674e-16 *** |
|  | 150 | 0.141 | 2.448e-11 *** | 0.131 | 5.906e-10 *** |
| V1/V4 | 20 | -0.013 | 0.392 ns | -0.004 | 0.779 ns |
|  | 150 | -0.032 | 0.032 * | -0.025 | 0.091 • |

By comparison, the pattern of correlations between V1 and V4 (Fig.5E&F) were therefore different from within-area correlations in two important ways. First, there was no measurable relationship between *r_CCG_*(20ms) and *r_signal_*. Second, at longer time-scales, a weak relationship emerged, but it was negative (see Table 2). That is, V1/V4 pairs tended to have opposite noise and signal correlations (like in Fig.3D–F), unlike the pairs from within the same area (*e.g*. Fig.3A–C). The pattern of correlations between V1 and V4 suggests that a different principle governs their relationships. These between-area correlations are also more evident at longer time-scales.

A concern might be that some of these relationships could be driven by the fact that a dynamic RDS stimulus with the same properties will vary subtly from trial to trial in the exact positioning and sequencing of the dots. Such variation could conceivably induce a local correlation in firing due to similar responses to the details of the dot sequence rather than the same binocular depth. One way to address this is to construct pseudo-random sequences of dot presentation that are in fact identical. This is achieved by creating so-called ‘frozen noise’, in which the same seed is given to the pseudo-random number generator on every trial, thereby creating identical sequences of dots on repeated trials. We performed some control experiments using frozen noise, comparing neuronal responses with and without resetting of the number seed. These experiments were performed with 100% binocular correlation, as this condition may be expected to give the strongest form of false association between binocular depth sensitivity and dot sequence. As with previous uses of this approach (Britten, Newsome, Shadlen, Celebrini, & Movshon, 1996), we found that the details of the dot sequence did not matter substantially. Specifically, the information about binocular depth was not artificially inflated when frozen noise, rather than new random sequences, was used to generate the RDS stimuli.

### Empirical evidence of differential noise correlations

A property of information-limiting correlations is that they limit discrimination thresholds in the neighbourhood of a specific stimulus value (Moreno-Bote et al., 2014). For this reason, they are also called differential correlations, since the concern is with the component of noise that lies along the tuning curve contour corresponding to the stimulus change. Theory therefore predicts that measured noise correlations are the sum of two sources, one that limits the sensory threshold due to differential correlations and one that does not (Moreno-Bote et al., 2014).

The differential correlations that limit information about the sensory discrimination scale with the product of the first derivatives of the two tuning curves from the pair of neurons under consideration (Fig.3C). It is here that the specifics of our experimental design present an advantage, because every pair of neurons was tested across a range of binocular disparities, as the psychophysical task was being actively performed. Hence, the relationship between the size of the noise correlation and the value of the derivatives of the tuning curves could be measured at multiple points along the tuning curve for every neuron pair. Moreover, all of these measurements could be directly related to the behavioural performance of the task.

The other component of noise correlations does not limit information because, in principle, this source of noise could be eliminated by averaging or pooling across many neurons. Nonetheless, this source of noise does present a practical limitation because it may mask the differential correlations that need to be measured and characterized. We reasoned that the low variance measure of noise correlation, *r_CCG_*, would be more likely to reveal weak and rapid differential correlations, rather than *r_SC_*, which is noisier. Thus, we examined the relationship of *r_CCG_* at different integration times (*τ*) and the product of derivatives over a range of time scales.

For pairs of neurons from the same area (V1 or V4), noise correlations had a positive relationship with the product of derivatives (*f*′×*f*′, see equation 23), during both the Initial and Steady state phases of neuronal activity (Fig.6&S6). This relationship is shown for some specific values of time scale (*τ*), plotting *r_CCG_* against *f*′×*f*′ (V1 Fig6A&B; V4 Fig.6D&E,S6A&B; regression lines in orange). A continuous plot of the slope (Spearman correlation) between *r_CCG_* and *f*′×*f*′ across time scales (*τ*) is shown in Fig.6C for V1 and Fig.6F&S6C for V4; these continuous plots were used to select time scales for the illustrative plots (Fig.6A,B,D,E,S6A&B). The vertical arrows on the time-scale axes show the *τ* of the maximum regression slopes for V1 (Fig.6C) and V4 (Fig.6F&S6C). Statistical summaries for the plots in Fig.6 (A,B,D,E) are shown in Table 3. It is evident that the relationship between *r_CCG_* and *f*′×*f*′ is most prominent in a narrow range of time scales 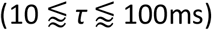.

**Figure 6.**
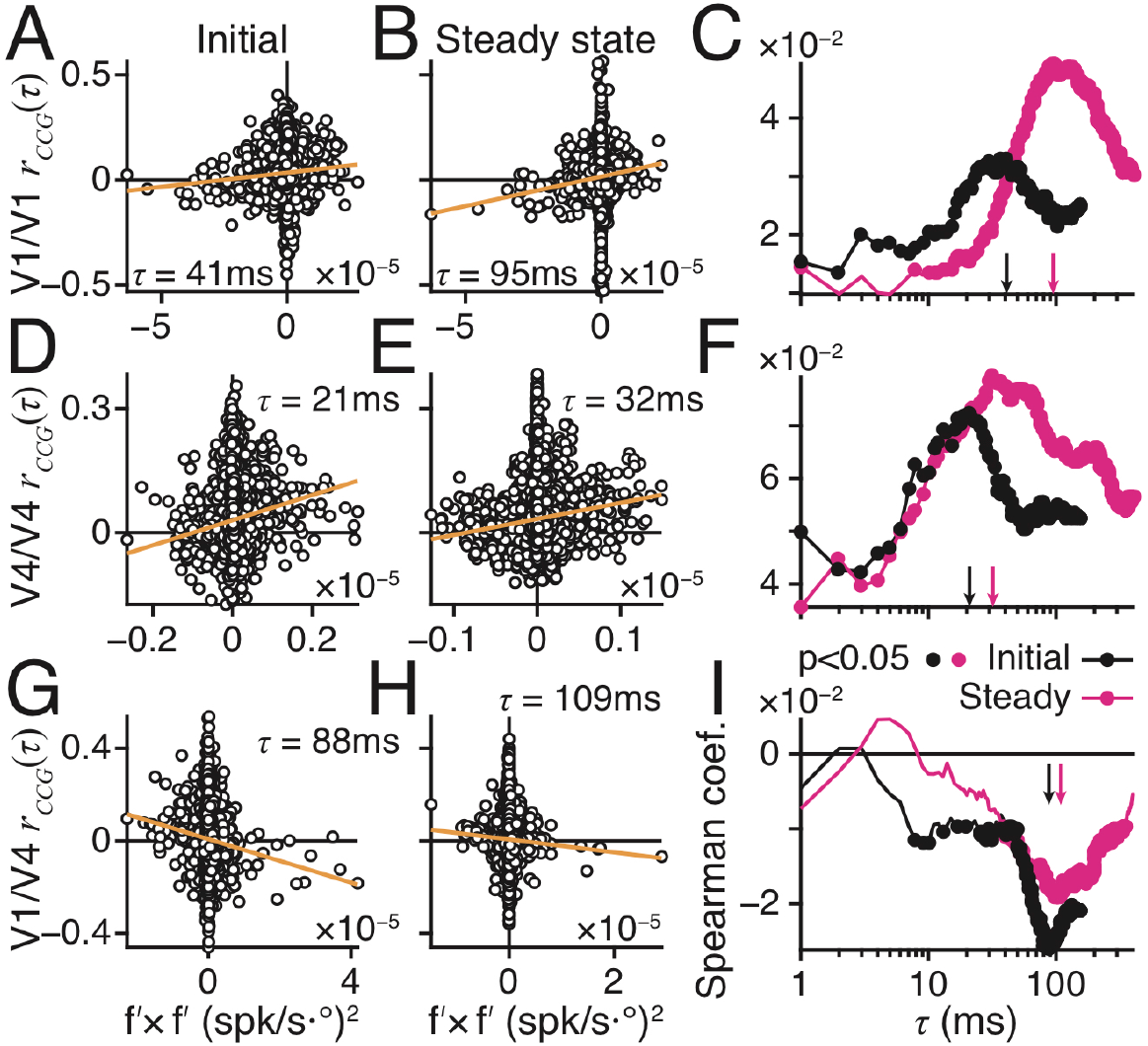
Empirical evidence of differential noise correlations. A) *r_CCG_*(*τ*) from V1 at *τ* = 41ms plotted over product of first derivatives of disparity tuning curves (*f*′×*f*′), for Initial response phase, N = 21,771 cases (2419 V1/V1 pairs at 9 disparities −0.1° to +0.1°). Best-fitting, least-squares regression line (orange) *r_CCG_*(τ) = *k* + *m*(*f*′ × *f*′) had positive slope (m): k = 3.452e–2; m = 1.244e–7. B) As A but for Steady state: *τ* = 95ms; k = 1.594e–2; m = 2.812e–7. C) Spearman correlation coefficient of *r_CCG_* and *f*′×*f*′ for Initial (black) and Steady state (magenta) responses, at all *τ* ≤ 400ms. Filled circles show p < 0.05 (two-tailed). Arrows show *τ* used in A (black) and B (magenta). D – F) Same as A – C but for V4/V4 pairs and using *τ* = 21ms for D and *τ* = 32ms for E, with N = 20,034. Positive regression slopes in D: k = 3.020e–2; m = 3.039e–6, and E: k = 3.330e–2; m = 3.891e–6. G – I) Same as A – C but for V1/V4 pairs and using *τ* = 88ms for G and *τ* = 109ms for H, with N = 41,382. Negative regression slope in G: k = 7.782e–3; m = −4.711e–7, and H: k = 6.354e–3; m = −2.804e–7.

**Table 3:** summary of relationship between r_CCG_ and f′×f′, accompanies Fig.6

| Cortical areas | Initial |  |  | Steady state |  |  |
| --- | --- | --- | --- | --- | --- | --- |
| | $\tau$ (ms) | Spearman $r$ | P-value | $\tau$ (ms) | Spearman $r$ | P-value |
| V1/V1 | 41 | 0.033 | 1.137e-6 *** | 95 | 0.049 | 3.423e-13*** |
| V4/V4 | 21 | 0.072 | 1.437e-24*** | 32 | 0.079 | 2.726e-29*** |
| V1/V4 | 88 | -0.026 | 1.023e-7*** | 109 | -0.019 | 9.298e-5 *** |

For pairs of neurons recorded simultaneously between V1 and V4, the result was unexpected (Fig.6G–I). Instead of a positive relationship, we observed a consistent negative relationship of *r_CCG_* and *f*′×*f*′ in the Initial and Steady state time periods, which peaked for a narrow range of *τ* close to 100ms (Fig6G,H,I). This negative relationship of *r_CCG_* and *f*′×*f*′ does not mean that the values of *r_CCG_* are themselves negative. On average, the values of *r_CCG_* and indeed of *r_SC_* are positive (see Fig.4). It is how these positive values of *r_CCG_* vary with the size and sign of *f*′×*f*′ that is of concern here, similar to the between-area relationship of *r_signal_* and *r_CCG_*, mentioned above (Fig 5 and Table 2).

### Differential noise correlations are attenuated at long time scales

The amount of information that can be encoded by a neuronal pool depends fundamentally on the number of different firing states that the pool can adopt. Thus, correlation among members of the pool reduces the capacity for information coding. For a sensory discrimination task, the impact of these correlations depends on the discrimination task that is to be performed. Differential correlations that mimic the change in neuronal response to the stimulus change that must be identified will result in a saturation of the information in a neuronal pool of a certain size. This is, differential correlations will follow the same principle identified by earlier studies, which concluded that threshold performance as a function of pool size is limited by correlation in the neuronal spike counts (Johnson et al., 1973; Shadlen et al., 1996).

Therefore, we measured the information about binocular depth that was available within neuronal pools of increasing size (Figure 7A&S7A). Information was assessed as linear Fisher information (equation 24)(Kanitscheider et al., 2015b), which measures how accurately the responses of the neurons in a pool can determine which binocular depth was presented. Fisher information is inversely related to the width of the likelihood function for estimating binocular depth. It therefore increases when the population response can better distinguish between different RDS depth values, and an increase of Fisher information is brought about from the addition of informative neurons or from changes in the pool’s correlations (Kohn et al., 2016). Empirically, information-limiting correlations can be removed from measured samples of neuronal population activity by shuffling the relationship between neuronal activity across multiple presentations. Shuffling decorrelates the population activity and is therefore is predicted to increase the Fisher information, if information-limiting correlations are present in the experimental measurements.

**Figure 7.**
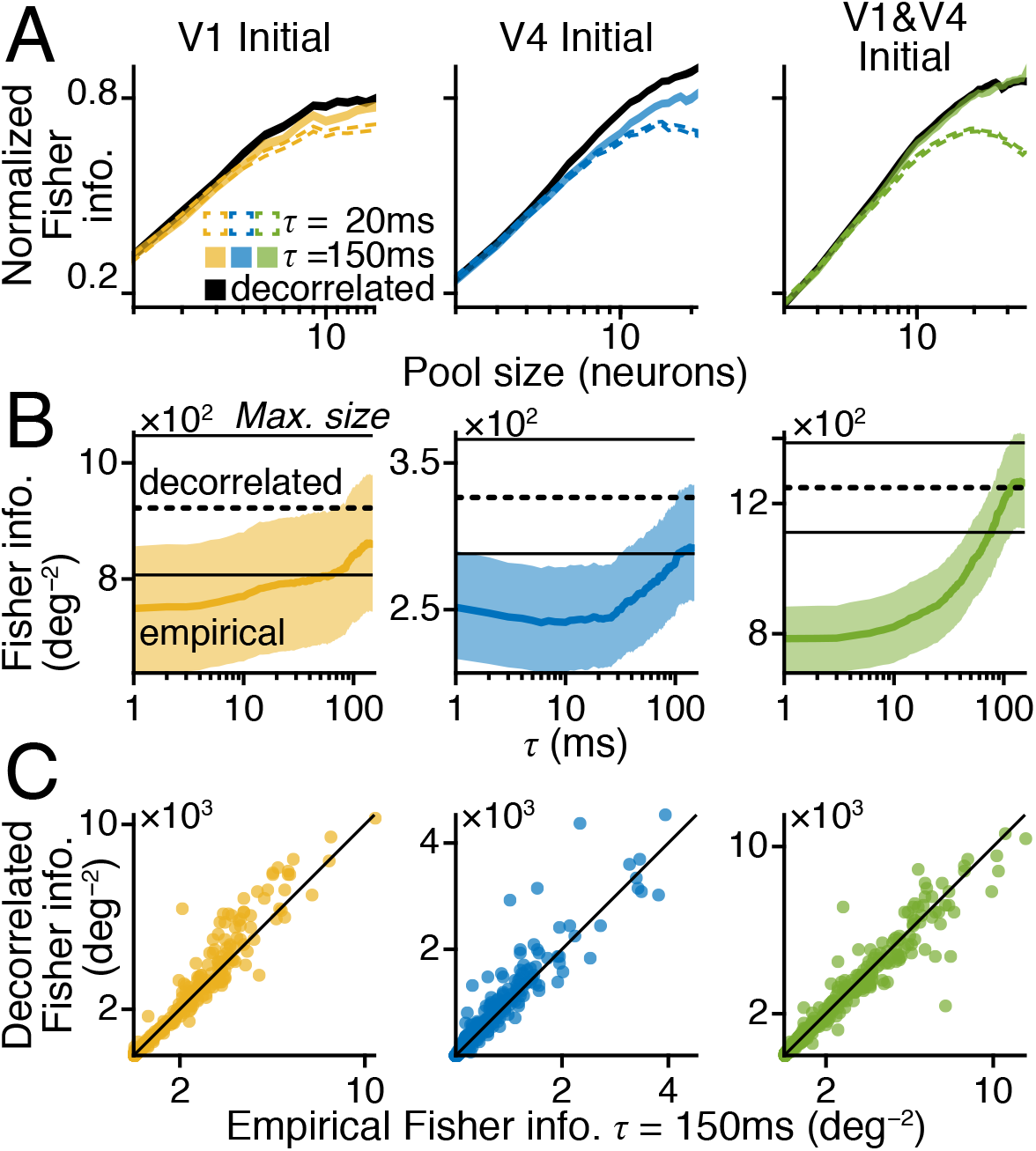
Differential noise correlations are attenuated at long time scales. A) Average ±SEM normalized bias-corrected linear Fisher information, over neuronal pool size. Information shown for rapid (*τ* =20ms, dotted), slow (*τ* =150ms, solid), and no (black) noise correlations, N = 780 (78 disparity pairs × 10 experiments). Units added to pools by decreasing order of Initial DMI.. B) Average Fisher information across *r_CCG_ τ*. Information is empirical (thick solid) or decorrelated (dashed). 95% bootstrap CI. C) Decorrelated vs. empirical information at *τ* = 150ms. Max. pool sizes in B & C.

Using responses from the Initial phase of firing, we calculated the normalized and averaged bias-corrected linear Fisher information (equation 32) for pools of units from V1-only (gold, left), V4-only (blue, middle), and V1 mixed with V4 (green, left). Units were added to each pool from the most selective for disparity down to the least selective (Sanayei et al., 2018); thus, part of the apparent saturation is due to exhaustion of informative units rather than the effect of correlations. The range of pool sizes was limited to those observed in at least half of the experimental sessions. The Fisher information available from experimental recordings was compared against the information within an equivalent pool of decorrelated units (solid black curve), for the cases of rapid (*τ* = 20ms, dotted) and slow (*τ* = 150ms, solid coloured) noise correlations. For all pool types, we found that the empirical information at *τ* = 20ms began to diverge from the decorrelated information for pools of 10 units. In pools with only V1 or V4 units, there was a similar but weaker divergence when *τ* = 150ms. However, for the mixed V1 and V4 pool, the empirical information at *τ* = 150ms was closely aligned with the decorrelated information (Fig.7A, right panel).

To get more detail about the effect of time scale, we plotted the average, raw (not normalised) Fisher information of the Initial responses as a function of *τ* (Fig.7B&S7B), using the maximum available pool size for each pool type in each experimental session. The empirical information (solid coloured line) was compared against the decorrelated information (dashed, black horizontal line). For all pool types, the empirical information tended to increase with *τ*. But the empirical information converged on the decorrelated information only in the mixed V1 and V4 pool, at around *τ* = 100ms.

We confirmed this result by plotting the Fisher information after decorrelation against the empirical information for *τ* = 150ms (Fig.7C&S7C). The information after decorrelation was greater than empirical for the single area pools (V1: mean paired difference ‘PD’ = 66.117, paired-sample t-test *p* = 4.700e–6; V4: PD = 32.316, *p* = 1.745e–7), but there was no significant difference for the mixed pool (V1/V4: PD= −20.558, *p* = 0.183). Similar results were observed using Steady state responses (Fig.S7D-F), except that the empirical information never entirely converged on the decorrelated information over the range of time scales that we analysed.

We tested whether the increase in the mixed V1/V4 information might be due to its larger pool size, compared to the V1- or V4-only pools, rather than any differences in the structure of the noise correlations. This was tested by plotting the information of a mixed pool with 12 units against the sum of information from separate V1- and V4-only pools of size 6, each (Fig.S7G); as in Fig.7A, the most informative units were used. The average paired difference in information (Fig.S7H) revealed that the mixed pool had extra information, over a range of values for *τ*. This difference increased with *τ*; hence information increased in the mixed pool as its covariance structure changed.

### Differential correlations and receptive field separation

A proper analysis of this data set requires consideration of the relative positions of the receptive fields (RFs) recorded from V1 and V4. Due to the topographic placement of the chronic Utah arrays, the majority of V4 RFs were more eccentric than the V1 RFs. Furthermore, the average distance between V1 and V4 RFs was greater in one monkey (M135) than the other (M138). During our main experiments, we presented areas V1 and V4 with a single visual stimulus that extended across the RFs of both visual areas. Both set of RFs were therefore co-activated by a single RDS stimulus (Fig.2A).

Nonetheless, it could be argued that the presence of a correlation structure that is the opposite of normal information-limiting correlations is simply due to the RF separation rather than the fact that the recorded neurons were in different cortical areas. Further, the apparent cancellation of information-limiting noise might have been caused by the misalignment of V1 and V4 RFs rather than any genuine corrective mechanism. To address these concerns, we can formulate two, testable null hypotheses. First, when compared over a range of similar values of RF separation, there will be no difference in the structure of within-area and between-area correlations. Second, any measurable recovery of information by a population of V1 and V4 neurons should increase as the separation of V1 and V4 RFs increases.

To measure RF separation, we performed an independent quantification of the positions and sizes of the V1 and V4 RFs. RF position was mapped in separate experiments by placing a small random dot stereogram (RDS) at randomly selected positions on a grid of possible locations, while the animal maintained fixation; the RDS position changed between trials. The size of the RDS was fixed (2.5° diameter) and the binocular disparity was always 0°. Thus, we mapped the average multi-unit firing rate for each electrode across RDS positions. We quantified the RFs by least-squares fits of two-dimensional isotropic Gaussians to each RF map, with centre and standard deviation (SD) of the Gaussian taken as measures of the RF centre and size.

The Gaussian fits were reasonably accurate (median coefficient of determination *i.e*. r^2^; M135 V1 = 0.875, 95% boot CI [0.788, 0.881]; M135 V4 = 0.650, [0.517, 0.813]; M138 V1 = 0.966, [0.962, 0.968]; M138 V4 = 0.862, [0.739, 0.926]), exhibiting the expected, fundamental relationship between RF eccentricity and size (all RFs: Spearman correlation M135 = 0.829, p → 0; M138 = 0.634, p = 3.351e–9).

The RF separation of pairs of neurons was defined by computing the Euclidean distance between RF centers, in degrees of visual field. For neurons i and j (i ≠ j), RF separation was 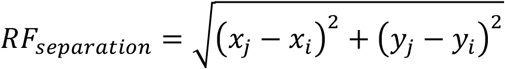 where (x_i_, y_i_) was the center of the Gaussian fit to the RF map of the i^th^ neuron, similarly for the *j*^th^ neuron.

Information-limiting noise correlations produce a positive relationship between measured noise correlations and the product of tuning curve derivatives (*f*′×*f*′); for brevity, we shall refer to this relationship as R_D_. Our main analysis revealed a positive R_D_ between pairs of neurons from the same cortical area, providing direct evidence of information-limiting noise correlations within both V1 and V4 for encoding binocular disparity. However, pairs of neurons from different cortical areas had, on average, a negative R_D_. Plotting R_D_ as a function of RF separation in Figure 8 shows that within-area correlations are generally positive, on average, whilst correlations between areas are predominantly negative – across the spatial separations in our sample. Neuron pairs were first binned (1 circle/bin) by their RF separation, with at least 100 pairs/bin (N bins; M135 V1/V1 = 3, V4/V4 = 9, V1/V4 = 10 (with outlier 11); M138 V1/V1 = 19 (20), V4/V4 = 11 (12), V1/V4 = 33); binning was done after discarding the 2% of pairs with the lowest DMI *i.e*. the least disparity selectivity. The R_D_ of each bin was quantified as the Spearman correlation of measured noise correlations (r_CCG_(150ms)) and *f*′×*f*′ of the bin’s pairs. Likewise, the average RF separation was found for each bin. There was no provable relationship between RF separation and differential correlations. If anything, our data indicate that all relationships, positive and negative, declined as the spatial separation of RF centres increased.

**Figure 8.**
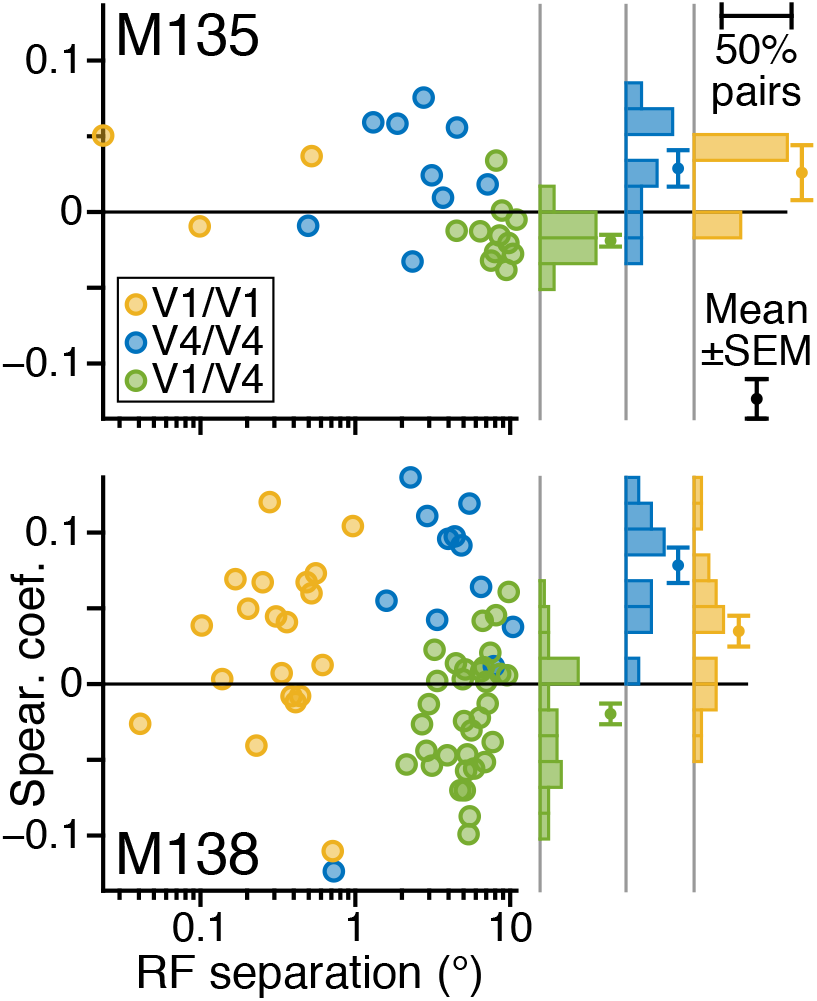
Differential correlations and receptive field separation. For each type of neuron pair, pairs were ordered by RF separation then grouped; at least 100 pairs to each group (N bins = N pairs / 100, rounded down). Spearman correlation of r_CCG_(150ms) versus *f*′×*f*′ for the pairs in each group, as a function of average log_10_ RF separation. Marginal histograms show distribution of Spearman correlation, counting groups.

We then asked how much linear Fisher information loss (equation 31) there was due to potential differential correlations (Fig.S8); this was taken to be the difference between empirical and decorrelated information levels, normalised by the expected root of the variance (equation 31). The amount of loss for population compositions was sensitive to the temporal scale (τ) of the measured noise correlations (Fig.S8A&B). The greatest information loss was seen at rapid (τ < 100ms) time scales. However, this information loss disappeared at longer time scales (τ > 100ms), suggesting an effective cancellation of information-limiting noise. To test the effect of RF separation on information loss, we plotted loss (Fig.S8C) across values of τ for two pools of neurons: 1) all V1 and V4 pairs with RF separations up to a criterion value of 7.99° (V1&Close-V4, green solid), and 2) the pool of V1 and V4 pairs with separations exceeding that criterion (V1&Distant-V4, dashed black). As before, we found strong evidence of information loss at small values of τ in both populations, and in both subjects. However, the V1&Close-V4 population was always the first to reach an average loss of zero. This effect was more prominent in M135, possibly due to the greater separation of the V1 and Distant-V4 RFs recorded from this cortex.

To better illustrate this point, we fixed τ at 120ms. At this temporal scale, the information loss of the V1&Close-V4 neurons was no different from zero in either M135 (average = +0.031, 95% Boot CI [-0.012, 0.075]; right-tail t-test p = 0.083) or M138 (−0.037 [-0.111, 0.043], p = 0.820). However, the loss of the V1&Distant-V4 neurons was significantly positive in M135 (0.125 [0.070, 0.182], p = 7.827e–6) and M138 (0.130 [0.036, 0.231], p = 0.005). A direct comparison between the two populations was possible, with the information loss of the V1&Distant-V4 population being significantly greater (right-tail paired-sample T-test p, M135 = 0.002, M138 = 2.731e–5). Our results suggest that the level of information loss increased with RF separation. They further suggest that information-limiting noise correlations are cancelled when the V1 and V4 RFs are close together.

In summary, our data support two conclusions. First, there is a categorical difference between within-area correlations and between-area correlations, regardless of RF separation. Second, information recovery between V1 and V4 is stronger when RF separation is smaller.

### Differential correlations and behavioural choice

In the final phase of the trial (Fig.1Aiv), the binocular depth instantaneously changed in one of the four RDS stimuli. This popout event revealed one out of the four RDS stimuli to be an oddball target. The monkey could get a reward by correctly choosing the target RDS with a saccade onto that stimulus. Rewards were given after the gaze fell reliably onto the target RDS (correct choice). Given that we recorded neuronal responses to just one of these four RDS locations, we divided the trials into four categories of outcome, specified by combinations of two factors: Target IN or OUT of the RF, and CORRECT or INCORRECT choice.

The association between behavioural choice and the neuronal state is plotted in Figure 9. The x-axis shows time relative to the pop-out event. As before, the state of the noise correlation structure of the recorded neuronal responses, R_D_, is summarized by the Spearman correlation between *r_CCG_*(*τ*) and the product of differentials *f*′×*f*′. Values of *τ* were 30ms for the V1/V1 and V4/V4 pairs, and 110ms for the V1/V4 pairs. Owing to differences between animals that emerged in early training, the two monkeys were exposed to either 1 second (M135) or 2 seconds (M138) of indistinguishable RDS stimuli during the Presentation phase (Fig.1Aiii), prior to the pop-out event. In a fraction of catch trials, no depth change occurred, to measure guess rates; these trials were discarded from this analysis.

**Figure 9.**
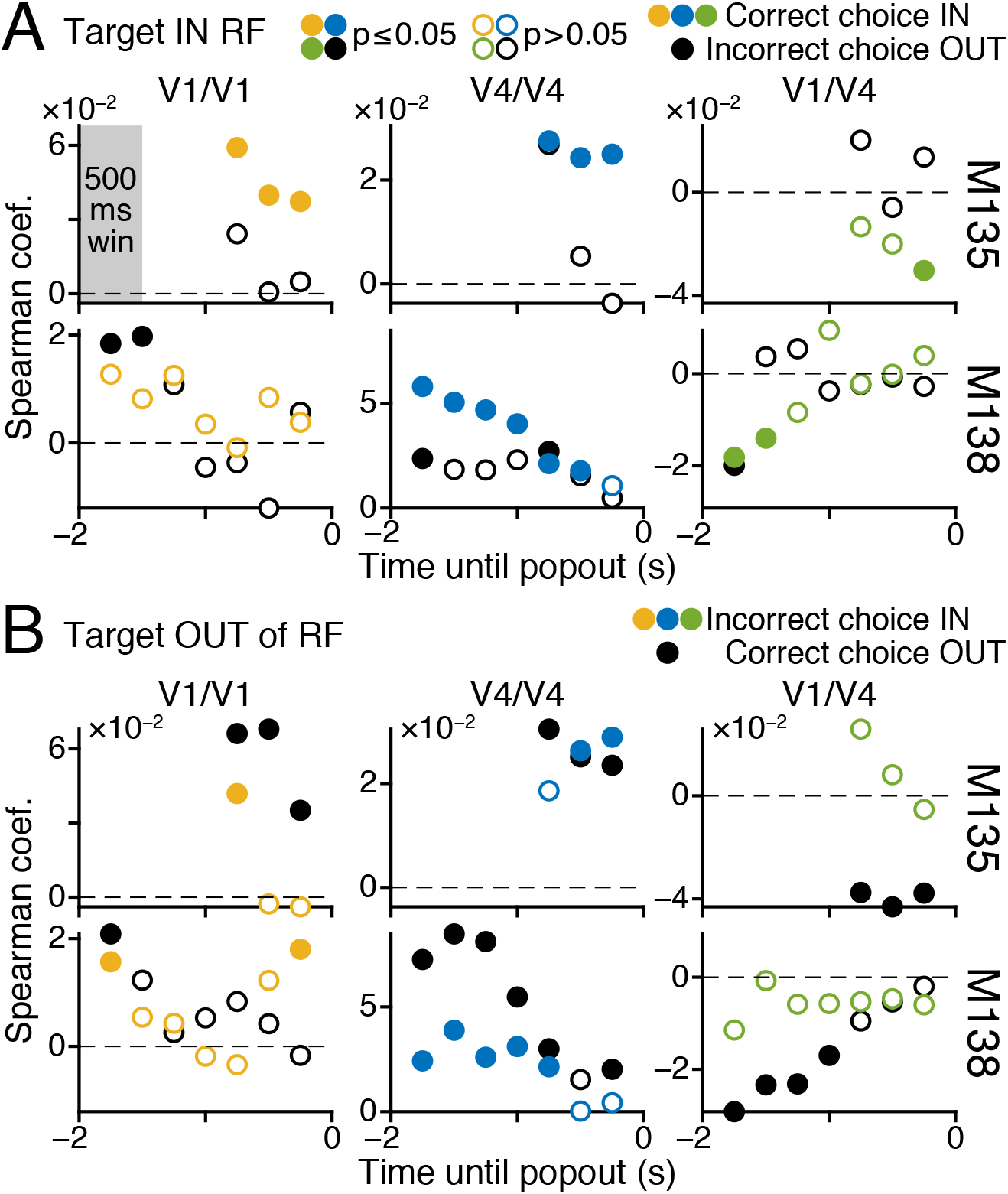
Effect of choice on direct evidence of differential correlation. Spearman correlation of r_CCG_(*τ*) versus *f*′×*f*′ (see Fig.6) in 500ms window (grey) at 250ms steps (circles, centred in each window) prior to popout event (Fig.1Aiv). For V1/V1 (left, *τ* = 30ms), V4/V4 (middle, *τ* = 30ms), and V1/V4 (right, *τ* = 110ms) pairs. RDS onset (Fig.Aiii) at −1 for M135), −2 for M138. Circle fill shows significance (two-tailed, p ≤ 0.05). Coloured circles show choices IN to RF, black show choices OUT of RF. A) Popout target IN the RF, choices IN are correct. B) Popout appears OUT of RF, choices IN are incorrect; incorrect choices out of RF not shown.

The data in Fig.9 show an association between the correctness of choice and the state of correlations, R_D_, prior to the behavioural decision. The effect is strongest for the within-area correlations of V4 (Fig.9A, middle panels, blue/black data points, fill shows two-tailed significance at 5% level). For both animals, correct choices onto the target over the RF are associated with stronger values of R_D_ in V4. This is not due to an alteration of correlations by the target stimulus, because the target is still unknown. Second, analysis of the opposite contingency (Fig 9B), when the target occurs outside of the RF, shows that these correlations are also stronger for correct choices. In this case, a correct choice means selecting a target outside of the RF and choosing the target over the RF is incorrect. For cases, where both the target and incorrect choices were out of the RF, the R_D_ measure of differential correlation was similar to when incorrect choices went into the RF (not shown). Therefore, this association in V4 is between correctness of choice and the measured state of differential correlations, regardless of target location. For the within-area correlations of V1 (Fig.9, left panels, gold/black data points), the same general trend is present, but the effects are much weaker and less consistent, especially in monkey M138.

For the between-area correlations of V1/V4 (Fig.9, right panels, green/black data points), an interesting pattern emerges. Previously in Figures 6&8, we demonstrated a negative Spearman correlation for the V1/V4 R_D_. This negative relationship is even stronger for correct choices as opposed to incorrect choices, most notably when the target is outside the RF (but note that target IN trial count is ≈25% of target OUT).

To quantify the significance of these changes, an all-ways ANOVA was performed on the normalised ratio of *r_CCG_* over *f*′×*f*′ (equation 35), separately per subject, testing the effects of time (window position), cortical area (V1 only, V4 only, V1/V4), and choice (correct or incorrect). The normalisation permitted the pooling of target IN and OUT trials, because both sets show a similar dependency on correctness. The effect of choice was significant in all but one condition (M135, V1 only, −500 to 0ms window, p = 0.979) when controlling for both time and cortical area (Tukey-Kramer multiple comparisons test, p < 0.001).

By themselves, these results do not necessarily confirm that differential correlations are stronger in magnitude when correct choices are made (that is, more positive for within-area correlations or more negative for between-area correlations). Another interpretation is that correct performance is associated with a suppression of general, non-differential correlations within the neuronal pool, and that this suppression allows a stronger coupling between information-limiting correlations and task performance.

More direct evidence of the potential importance of differential correlations is presented in Figure 10. Here, the goal is to obtain a unidimensional measure of the differential correlation in a neural population on each trial, so that its trial-to-trial relationship with perceptual choice could be quantified. The focus is upon the change in the V1 and V4 population response that is caused by the popout stimulus. By projecting the population response onto the optimal axis for discriminating baseline vs. popout disparity (equation 36), we reduce the N-dimensional firing rate down to a 1D estimate of the popout disparity. This is done by taking a weighted, linear sum of the firing rates, using weights given by w~Σ^−1^ΔF, for population covariance Σ and expected size and sign of response to the popout, ΔF; weights were optimised according to the baseline and popout disparity on each trial. Owing to the Σ^−1^ term, the optimal axis can mitigate non-differential noise correlations, but any differential correlations in the population response are necessarily projected onto the optimal axis.

**Figure 10.**
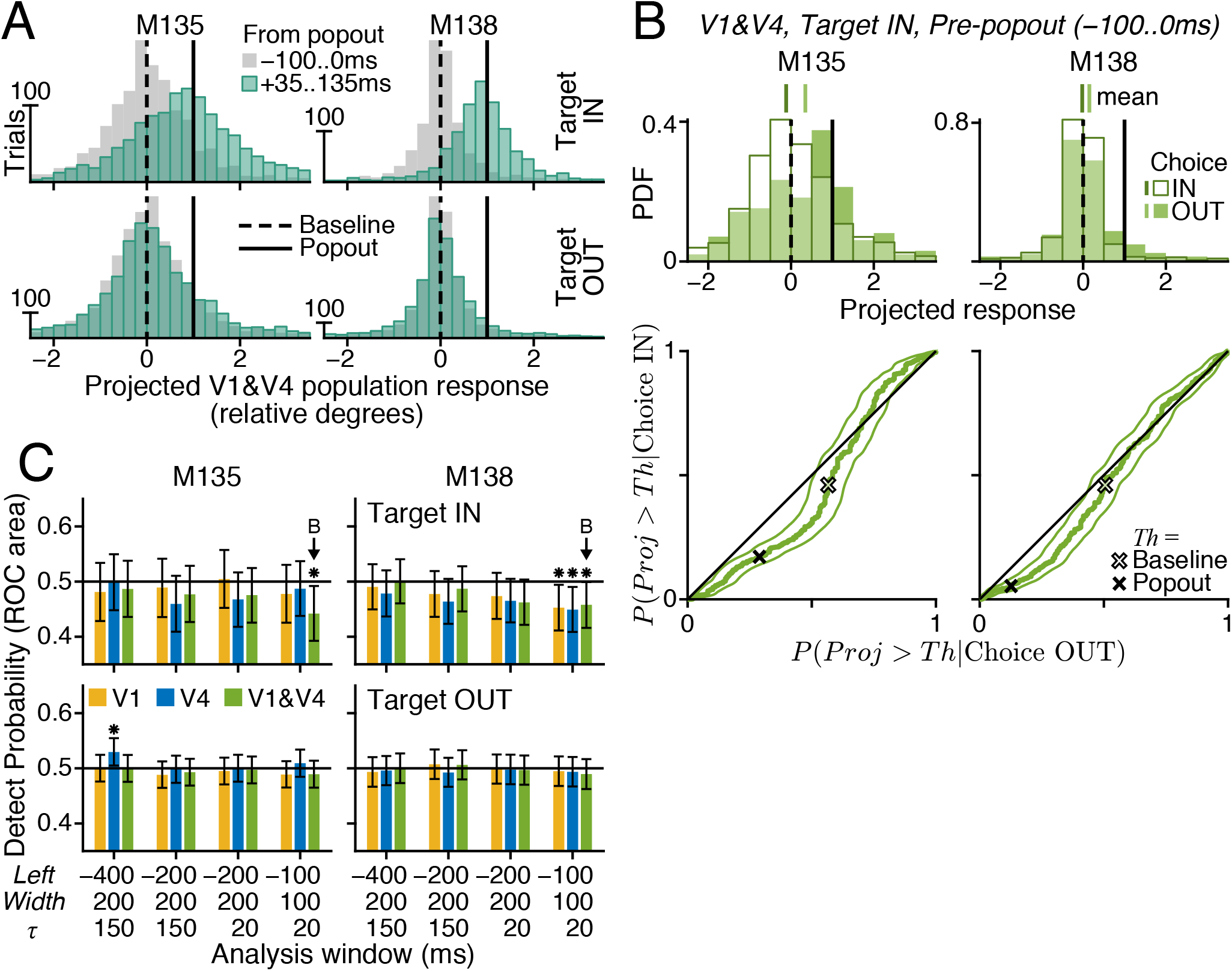
Differential correlation predicts choice. N-dimensional firing rates of simultaneous population responses were linearly projected onto the axis that optimally discriminated baseline from popout responses. A) Example projected responses for combined V1 and V4 populations. Expected value of 0 for response to baseline disparity (dashed), and +1 for popout (solid). Target RDS appeared IN (top panels) or OUT (bottom) of the RF. Responses prior to (−100 to 0ms, grey) or after start (+35 to +135ms, teal) of popout event. B) Example ROC curves (bottom, green) using pre-popout projections from A (top, grey). Black line: chance association. Crosses show probability values equal to expected baseline (open) or popout (filled) response. Top shows distributions grouped by choice IN (empty) or OUT (filled) of RF, black vertical lines as in A, green vertical lines show averages. C) Detect Probability for populations with V1 units only (gold), V4 units only (blue), or mixed V1 and V4 units (green), relative to chance (0.5, black horizontal) with target RDS IN (top) or OUT (bottom) of the RF. Population responses taken from analysis windows at different positions (left edge = −400, −200, or −100ms before popout) and width (200 or 100ms). Projection weights optimised for noise correlations at *r_CCG_* time scales *τ* = 20 or 150ms. * 95% CI different from chance. Arrows show DP for curves in B. All Error bars are 95% bootstrap CI (2e4 boot samples, percentile method). Neuron pairs *RF_separation_* ≤ 7.99°.

The values arising from this projection were normalised so that the expected response to the baseline disparity was 0, and the expected response to the popout disparity was always +1, regardless of whether it is actually an increase or a decrease. If pre-popout values for the projected response are near zero, this suggests that the population contained very little differential correlation and accurately encoded the baseline disparity. Values between 0 and +1 suggest the presence of differential noise that tilted the population closer to the firing pattern expected in response to the popout disparity, while values between −1 and 0 suggest the opposite. Hence the relative direction and magnitude of perturbations in disparity encoding could be estimated on each trial.

Fig.10A shows the distributions of projected responses from each animal from the combined population of all V1 and V4 units with *RF_separation_* ≤ 7.99° (see Fig.S8). When the analysis window is −100ms to 0ms (grey) before the popout event, then the distributions peak around 0, regardless of the upcoming target location. After the start of the popout event (+35ms to +135ms window, teal), the distributions shift towards +1, for the case when the popout target was IN the RF (top). For the other cases, when the target was OUT of the RF, then the pre- and post-popout projection distributions were indistinguishable (bottom). Thus, the normalised projection succeeded in representing the combination of signal and noise in encoding the RDS disparity.

We asked whether perturbations in this projected response that occur just prior to the arrival of the popout stimulus might affect the ability of the animals to detect the upcoming change in the visual stimulus. We quantified this by computing a version of Detect Probability (DP) (Cook & Maunsell, 2002) to measure the covariation of the normalised projection with the perceptual choice. Projected responses from the time period −100ms-0ms were now grouped according to behavioural choice (Fig.10B top, Choice IN (open) and OUT (filled) of the RF). These distributions, categorized by future choice, allow the computation of receiver operating characteristic (ROC) curves (Fig.10B bottom, green), where the choice IN rate is plotted on the vertical axis of the ROC curve and choice OUT is on the horizontal axis. The DP is the area under this ROC curve (equation 40), such that a DP > 0.5 indicates that larger, more positive perturbations preceded choices IN to the RF, while DP < 0.5 suggests the opposite and DP = 0.5 indicates no effect on detection. Fig 10B shows that both ROC curves dip below the chance line (black). This means that positive perturbations that mimic the arrival of the target stimulus and occur just beforehand have a negative effect on detection rates for the target.

It is interesting to note that the portions of the probability distributions of the projected response that are most relevant for this effect on behaviour lie exactly within the bounds that are also relevant for the neural detection of the target stimulus. The relevant positions are marked as the baseline (empty cross) and popout (filled cross) disparities and are simply transposed from the vertical lines in Fig 10A. The perturbations in the pre-popout period have their greatest impact on behaviour, producing a DP < 0.5, when they occur within these boundaries.

In Fig.10C, we computed DP for populations of units, with *RF_separation_* ≤ 7.99°, for V1 only (gold), V4 only (blue), or mixed V1&V4 units (green), using responses measured from a range of pre-popout windows (−400 to −200, −200 to 0, −100 to 0ms), with weights optimised using either fast (*τ* = 20ms) or slow (*τ* = 150ms) noise correlations; values were computed separately for trials with the target IN the RF (top) or OUT (bottom). We found that the only difference in DP that was consistent across subjects was in the combined V1 and V4 responses just prior (−100 to 0ms) to the appearance of the popout target IN the RF (arrows). Notably, the relevant temporal scale for these noise correlations was fast, at 20ms. This suggests that rapid, pre-popout differential correlations that spanned both V1 and V4 are detrimental to detecting the change in disparity.

This is a key prediction of the role for differential correlations in limiting the detection of the stimulus. When the state of the network before the arrival of popout is such that it more closely resembles the state of the network after the popout stimulus arrives, then this is detrimental to perception of the disparity change. On trials when the fluctuations in network state place it further away from the neural response to the popout, then detection of the popout target is less impaired. In line with Figs.6,7,&S8, our results suggest that it is the rapid differential correlations that do indeed degrade behavioural detection. Slower correlations prior to the popout event do not appear to impair detection, suggesting that integrative processes in the visual cortex may have enough time to attenuate rapid differential fluctuations.

### A model V4 output neuron can attenuate differential noise

Here we consider how the joint actions of V1 and V4 could preserve information transmission in the presence of information-limiting correlations within each individual cortical area. The finding of pairs of V1 and V4 units that had opposite disparity preferences and positive noise correlations (*e.g*. Fig.3D–F, Fig.S3 left) provides evidence for one route to cancelling the correlated noise without reducing the sensory signal (Abbott & Dayan, 1999; Averbeck et al., 2006). This phenomenon has been observed experimentally in the second order areas of the somatosensory system (Romo, Hernandez, Zainos, & Salinas, 2003).

To investigate this further, we built a simple model of a V4 output neuron (Figure 11, equations 37–39), of a type that might project to downstream areas. The model had two afferent neurons (Fig.11A), a neuron that projects from V1 onto the V4 output neuron and an intrinsic, local-circuit neuron from within V4 that also connects to the V4 output neuron. The details of this could be varied without a loss of generality, but importantly this represents a simple feedforward system that can explain many of our results.

**Figure 11.**
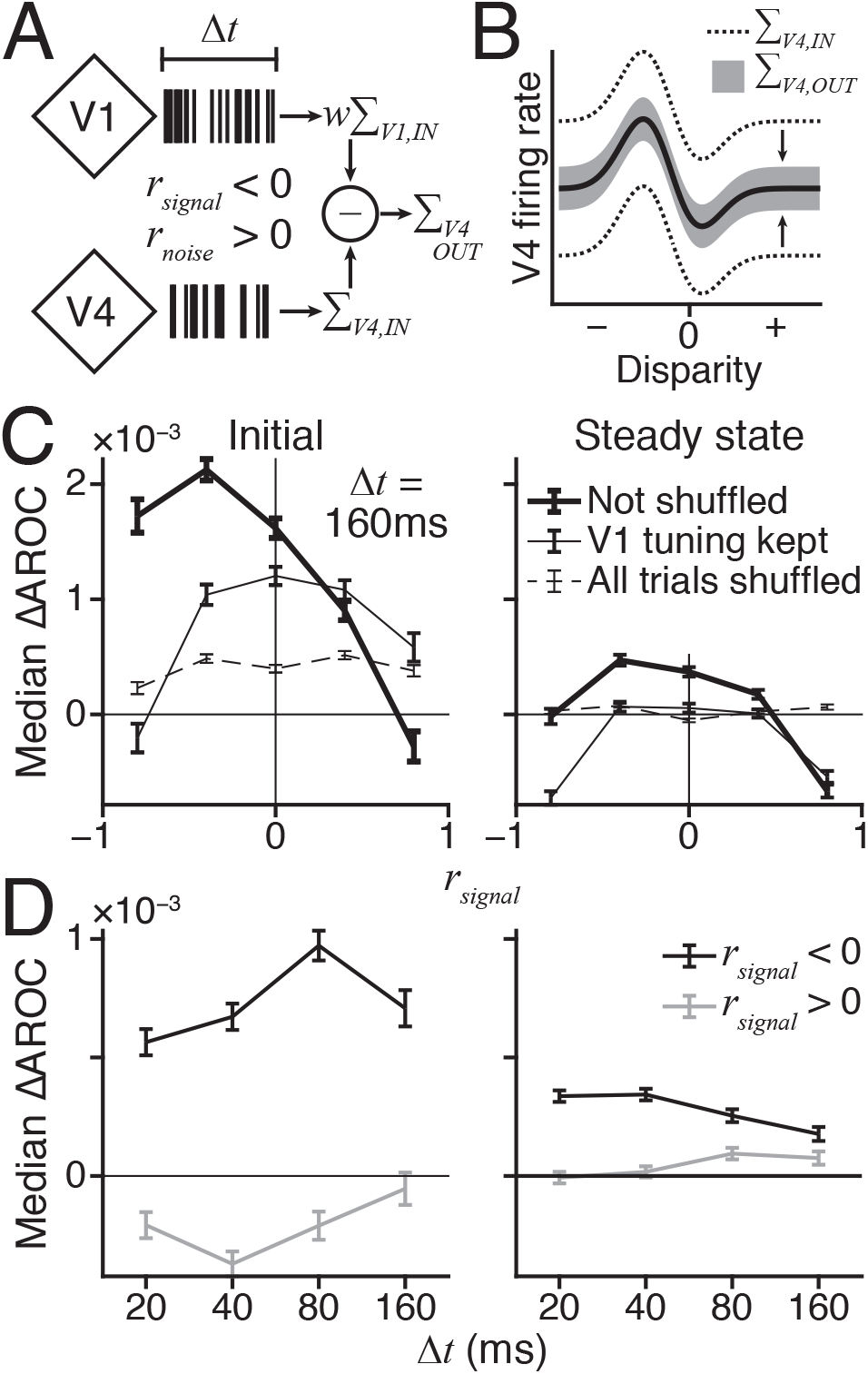
A model V4 output neuron attenuates differential noise. A) Diagram of model V4 output neuron. Diamonds, afferent neurons. Rasters, afferent spikes in Δ*t* ms window. Weighted V1 count (*w*∑_V1,IN_) subtracted from V4 count (∑_V4,IN_) gets V4 output count (∑_V4,OUT_). B) Disparity tuning prediction, afferent V1/V4 *r_signal_* < 0 and *r_noise_* > 0. V4 input variance (∑_V4,IN_, dotted). Model V4 output variance (∑_V4,OUT_, grey). C) Average change in AROC across *r_signal_* of afferent V1/V4 pairs, Δ*t* = 160ms. V1 responses shuffled across trials to break V1/V4 correlations but not disparity selectivity (V1 tuning kept, thin), to break correlations and V1 tuning (All trials, dashed), or not shuffled (solid). N = 358,644 (78 baseline disparity pairs × V1/V4 pairs). Each bin has 10% of data. D) Average difference in AROC over Δ*t*, V4 model output AROC with no shuffling minus shuffled AROC with preserved V1 tuning. Afferent V1/V4 *r_signal_* < −0.5 (black) and *r_signal_* > +0.5 (grey). N = 4598 model V4 output neurons, one per recorded V1/V4 pair. C&D) Initial data on right, Steady state on left. SEM error bars.

Correlated noise was removed by subtracting the weighted V1 responses from the input V4 responses, each input integrated across a Δ*t* ms window, to produce the V4 output responses. Each V1 and V4 pair in our dataset was used to simulate the responses of a model V4 output neuron, whose disparity selectivity was then measured. The prediction is that the V1/V4 input pairs with positive noise correlation (*r_noise_* > 0) and negative signal correlation (*r_signal_* < 0) should yield V4 output neurons with better disparity selectivity (Fig.11B, V4 input variance, dashed, output V4 variance, grey).

We test this circuitry directly with empirical data by measuring the disparity selectivity of each simulated V4 unit, using the area under the receiver operating characteristic curve (AROC, equation 40) as a measure of neuronal sensitivity. Focusing on the Initial analysis window, the difference in AROC (equation 41) between the V4 model output and input (Fig.11C left, thick solid, Not shuffled) confirmed the prediction above, where positive differences indicate model V4 output neurons with better disparity sensitivity.

As stated, a pair could improve its stimulus sensitivity either by changing its correlation structure, or by producing a larger modulation of firing in response to stimulus changes. Hence, it is possible that the superior performance of the model V4 output neuron was trivially due to the combined signalling of the V1 and V4 input neurons, rather than any cancellation in correlated noise. We tested this by recomputing the model V4 outputs after shuffling the afferent V1 responses between trials such that the V1 unit maintained its stimulus sensitivity, but the noise correlations were destroyed (thin solid, V1 tuning kept). In this scenario, the model V4 neuron can only improve its sensitivity through combined input signals, but not through cancelling out the correlated noise of its afferents.

Model V4 neurons had better stimulus sensitivity than the afferent V4 neuron, whether there was shuffling (thin solid) or not (thick solid). Yet, the model V4 neurons with no shuffling achieved a further increase in stimulus sensitivity in comparison to those with shuffling. This improvement in sensitivity occurred specifically when the afferent neurons had opposite disparity selectivity (*r_signal_* < −0.5, mean difference AROC = 2.961e–3, paired t-test *p* = 4.974e–112); in contrast, the model without shuffling suffered a reduction in sensitivity when the afferent neurons had similar disparity selectivity (*r_signal_* > +0.5, −7.894e–4, *p* = 3.269e–10). This is precisely the pattern that should be observed when the afferent V1 and V4 neurons have positive noise correlations (Abbott & Dayan, 1999; Averbeck et al., 2006). Additional stimulus sensitivity is available by cancelling out the noise correlations of afferent neurons with opposite stimulus selectivity.

As differential correlations seemed to be reduced by slower, V1 to V4 correlations (Fig.7&S8), we asked if the simulated V4 output neuron would improve its performance with a wider integration window. Thus, we evaluated the Initial V4 output responses over a set of Δ*t* values. For each Δ*t*, the un-shuffled model V4 sensitivity (Fig.11D, left) was subtracted by the shuffled model V4 sensitivity; again, this tests for changes in sensitivity due to the cancellation of afferent noise correlations. When the input V1/V4 pair had *r_signal_* < −0.5, there was a significant tendency for sensitivity to increase as Δ*t* increases (Δ*t* = 20ms vs 160ms, mean paired difference = 1.540e–3, *p* = 9.490e–25).

For the Steady state responses, the AROC measure detected a weak improvement (Fig.11C, right) when the input pair had opposite disparity preferences (Not shuffled vs V1 tuning kept, mean paired difference = 4.9181e–4, paired t-test p = 3.1995e–26). No improvement in selectivity was found with increasing Δ*t* (Fig.D right, Δ*t* = 100ms vs 800ms, mean paired difference = −3.902e–4, *p* = 7.986e–16).

## DISCUSSION

Cortical areas V1 and V4 contain noise correlations that could limit the information that those areas convey about stereo depth. This noise limits information chiefly at short time scales, as the animal is discriminating depth. Unexpectedly, noise correlations between the two areas were found to have a different structure at longer time scales, which could reflect the brain’s capacity to limit correlations that might become harmful to perceptual performance. We show that the state of short-time course noise correlations has a direct association with the animal’s performance, demonstrating a critical piece of evidence that the structure of correlations has a direct impact on behavioural decisions and sensory performance.

### Detecting and interpreting differential correlations

Empirical evidence of differential correlations has rarely been reported, due in part to the prediction that they are so small (Moreno-Bote et al., 2014) that thousands of neurons and thousands of trials are needed to detect them (Kanitscheider et al., 2015b; Kohn et al., 2016).

But the magnitude of noise correlations is dynamic, and changes with a number of factors (Marlene R. Cohen & Kohn, 2011), including state of consciousness (Ecker et al., 2014), stimulus presentation (Churchland et al., 2010), perceptual grouping (Bondy et al., 2018; M. R. Cohen & Newsome, 2008; Poort & Roelfsema, 2009; Wasmuht, Parker, & Krug, 2019), attention (M. R. Cohen & Newsome, 2009; Mitchell, Sundberg, & Reynolds, 2009; D. A. Ruff & Cohen, 2014; Verhoef & Maunsell, 2017), and perceptual learning (Ni et al., 2018; Sanayei et al., 2018). Our experiment brought many of these global factors under control by using awake subjects that performed a demanding perceptual task. Indeed, we presented evidence (Figs.9&10) that the animal’s fluctuations in perceptual performance may reflect the impact of differential correlations on vision perception.

Another advance we have made in detecting differential correlations is in how they are measured. Spike count correlation (*r_SC_*) is often used despite the fact that correlations vary over temporal scales (Bair et al., 2001; Denfield, Ecker, Shinn, Bethge, & Tolias, 2018; Kohn & Smith, 2005; M. A. Smith & Kohn, 2008; M. A. Smith & Sommer, 2013). If differential correlations are rapid, then they could be masked by the slower correlations that are captured by *r_SC_*. By using the *r_CCG_* measure introduced by Bair et al. (2001), we found evidence of differential correlations operating at tens of milliseconds. This is much shorter than the width of the analysis windows that have typically been used to measure *r_SC_* (Marlene R. Cohen & Kohn, 2011).

A complication is that the time scale of noise correlation is dynamic, and can change within an area due to stimulation or behavioural state. This can be seen by comparing our results with previous studies of V1 (Kohn & Smith, 2005; M. A. Smith & Kohn, 2008) and V4 (M. A. Smith & Sommer, 2013) that used *r_CCG_*. They found noise correlations that saturated around 100ms during stimulation, in both areas; but the subjects were either anaesthetized (Kohn & Smith, 2005; M. A. Smith & Kohn, 2008) or passively fixating (M. A. Smith & Sommer, 2013).

We saw a similar *r_CCG_* profile in the spontaneous responses of V1; but with visual stimulation presented under a task-relevant contingency, *r_CCG_* saturated at 20ms. In V4, saturation was also near 20 ms – before and during stimulation. Our results here are more closely compatible with other cases in which *r_CCG_* also saturates at tens of ms in the dorsal area V5/MT when monkeys perform a perceptual task (Bair et al., 2001; Wasmuht et al., 2019). Thus, the rapid, differential correlations that we observed here may have only been readily detectable during the performance of a perceptual task.

### Recording depth and topography

We used 1mm electrodes, with the result that our recordings were likely from the supragranular layers near the layer 3-4 border, from a mixture of neurons that projected nearby or to other cortical areas (Harris & Mrsic-Flogel, 2013). The correlations may have carried a stronger differential component if many neurons projected locally. This stems from the theory that differential correlations build up between neurons that share some of the same noisy afferents (Kanitscheider et al., 2015b). Given the functional heterogeneity across layers in V1 and V4 (Buffalo, Fries, Landman, Buschman, & Desimone, 2011; Denfield et al., 2018; Nandy, Nassi, & Reynolds, 2017; Pettine, Steinmetz, & Moore, 2019), cross-laminar recordings will be required to resolve this issue.

Some of the signal from V1 will have been relayed to V4 by other areas; but there are direct axon projections from V1 to V4 and from V4 to V1 (Ungerleider, Galkin, Desimone, & Gattass, 2008). Although we did co-activate the V1 and V4 neurons with a single stimulus, the measured V1 RFs were not perfectly coincident with those of V4. Hence, some of the excitatory drive for the V4 neurons presumably came from other, unmeasured V1 neurons. Importantly, we saw qualitatively similar V4 correlations in a set of control experiments with the RDS centred on the aggregate V4 RF (see Figs.S1&S2, S4–S7).

In V1 and V4, noise correlations span a range of cortical (*i.e*. retinotopic) distances when neurons respond to the same stimulus (M. R. Cohen & Newsome, 2009; Poort & Roelfsema, 2009; M. A. Smith & Kohn, 2008; M. A. Smith & Sommer, 2013) – which we observed (mean *r_CCG_*(20ms); V1 pairs < 1.44mm apart = 4.987e–2; V1 > 1.44mm = 1.805e–2; V4 < 1.44mm = 3.416e–2; V4 > 1.44mm = 2.279e–2; all t-test *p* → 0; median distance = 1.44mm). Moreover, noise correlations in both areas increase when neurons respond to a common stimulus amongst an array of stimuli (Poort & Roelfsema, 2009; D. A. Ruff & Cohen, 2014). Thus, it is likely that the correlations of perceptually grouped neurons vary systematically across the cortex. We suggest that by stimulating V1 and V4 with the same object that we increased the likelihood of the visual system having grouped them into a perceptual pool.

### Mechanisms of noise removal

In a sense, our results are puzzling. Neuronal correlations in V1 and V4 show positive relationships between the neuron pairs with similar stimulus tuning and tuning curve slope (differential correlations). This kind of pattern is reported for motion in V5/MT and orientation in V1. With simple feedforward connections, one ought to observe the same relationship between V1 and V4. However, the general pattern of V1/V4 correlations is inverted (Fig.5E&F,6I,8).

Noise correlations like that can be removed linearly without losing information (Abbott & Dayan, 1999; Averbeck et al., 2006). Thus, our simple model (Fig.11) suggests that the combined action of V1 and V4 attenuates the local correlations through linear operations. However, differential correlations cannot be linearly removed by later processing (Kanitscheider et al., 2015b; Moreno-Bote et al., 2014; Pitkow et al., 2015). One way of resolving this discrepancy rests on our result that the covariance structure between areas is not static, but evolves across temporal scales. Notably, the V1/V4 pattern emerged at slower time scales than the rapid differential correlations within each area.

Related results come from MT (M. R. Cohen & Newsome, 2008) and V4 (D. A. Ruff & Cohen, 2014) in which positive noise correlations strengthened between neurons with opposite tuning (but see Bondy et al., 2018). In V1, positive correlations that increase the information in linearly-pooled responses can appear between neurons that respond to different stimuli (Poort & Roelfsema, 2009). Lastly, to change the V1 to V4 correlation structure in a trial would require the kind of fast and highly specific adaptations that are observed between them in attention tasks (Grothe, Neitzel, Mandon, & Kreiter, 2012; Rohenkohl, Bosman, & Fries, 2018).

Adaptable covariance may imply an underlying, non-linear computation. But what could be biologically plausible? A spike-count variance code could overcome correlations if correlated inputs were high-pass filtered in a spatial domain (*e.g*. across neurons) followed by a quadratic non-linearity to compute variance (Shamir & Sompolinsky, 2004). Another potential choice is divisive normalisation (Carandini & Heeger, 2012), which can model the structure of correlations between areas V1 and MT (Douglas A. Ruff & Cohen, 2016). Divisive normalisation might decorrelate responses in two ways (see Eq. 3.1 in Tripp, 2012). First, spike count variance can be saturated by the correlated variance of other neurons, attenuating correlations in the normalised response. Second, the normalised responses of correlated units can move in opposite directions to their raw values, decorrelating them. One mechanism need not exclude others, and the brain might profit from the parallel action of several computations.

### Sources of differential correlations

Present analysis suggests both bottom-up (Kanitscheider et al., 2015b) and top-down (Ecker, Denfield, Bethge, & Tolias, 2016) origins of differential correlations. In the former, stimulus noise is sent through diverging projections to downstream areas; differential correlations then build up between neurons with common noisy inputs (Kanitscheider et al., 2015b), especially if sub-optimal computations are used (Beck, Ma, Pitkow, Latham, & Pouget, 2012). In the latter, noise in the selection of an attended feature value can lead to a gain profile that shifts position relative to the stimulus preferences of a population of neurons – inducing differential correlations (Ecker et al., 2016).

Our results invite a compromise. Both sources are present, but take effect at different moments. The initial burst of firing is a bottom-up event in which information is most reduced by rapid noise correlations, which are unlikely to have a top-down origin (Denfield et al., 2018). In our data, only the Initial responses could reach the decorrelated information level, at a temporal scale of 100ms. The Steady state information did increase at wider temporal scales, yet failed to hit the decorrelated level within an 800ms time scale. One interpretation is that rapid differential correlations in the Initial response had a bottom-up source, followed by slow differential correlations with a top-down source that could not be locally attenuated. This fits well with the common reports of top-down or contextual effects that follow the initial response of visual neurons to a stimulus (Roelfsema, Lamme, & Spekreijse, 1998) (Pack & Born, 2001) (M. R. Cohen & Newsome, 2009; Herrero, Gieselmann, Sanayei, & Thiele, 2013; Nienborg & Cumming, 2009; Poort et al., 2012; J. E. T. Smith et al., 2015).

### Neural pooling and perception

The statistical relationship between a pool of sensory neurons and perceptual behaviour is connected to the noise correlations within the pool (Haefner, Gerwinn, Macke, & Bethge, 2013; Nienborg, Cohen, & Cumming, 2012; Andrew J Parker, 2013; Pitkow et al., 2015; Shadlen et al., 1996). That relationship may evolve over time, with a sensory pool driving a brief, bottom-up effect on behavioural choice, followed afterwards by a top-down reflection of choice bias (J.E.T. Smith, Masse, Zhan, & Cook, 2012). These effects are difficult to separate in many experiments, which employ long stimulus viewing times (Britten et al., 1996; Dodd, Krug, Cumming, & Parker, 2001; Nienborg & Cumming, 2009; Pitkow et al., 2015; Poort et al., 2012; Yu & Gu, 2018).

However, a subject may base perceptual decisions on only a few hundred ms of viewing in natural conditions (M. R. Cohen & Maunsell, 2009). Hence, studies using mixtures of reaction time tasks and short-duration stimuli often observe that the transient response is accounted for by a small pool of neurons (Ghose & Harrison, 2009; Harrison, Weiner, & Ghose, 2013; Krause & Ghose, 2018; K. F. Weiner & Ghose, 2015) with a bottom-up link to behaviour (Price & Born, 2010; Jackson E. T. Smith, Zhan, an, & Cook, 2011; Katherine F. Weiner & Ghose, 2014). Our results suggest that the transient response is a high-fidelity signal that could be decoded from a small pool of neurons. Later, top-down signals may arrive that impose slow and widespread differential correlations in a larger pool, thus injecting choice correlations into neurons that are never read out (M. R. Cohen & Newsome, 2009; Nienborg & Cumming, 2010; Nienborg & Cumming, 2009; Pitkow et al., 2015; Yu & Gu, 2018).

Although the associations between behavioural decisions and network state were stronger in one monkey than the other, the timing of these changes of state, in terms of differential correlations, supports this interpretation. Rapid fluctuations in the state of the network that mimic the change of state that occurs upon the presentation of the discrimination stimulus are detrimental to stimulus detection. This is a fundamental prediction of the theoretical role of differential correlations (Pouget et al., 2003) that we have been able to validate with these experimental measures.

### Correlations and the organizational structure of the neocortex

Our results lead us to propose a general framework for thinking about the effect of noise correlations on neuronal processing. First, within-area noise correlations that potentially limit sensory discrimination are a by-product of local information-processing (Harris & Mrsic-Flogel, 2013; Kanitscheider et al., 2015b). Second, we suggest that the signalling between areas acts to remove the harmful consequences of these local, intrinsic correlations. The removal process may use at least two strategies: one is the connectivity between cortical areas (*e.g*. Figure 8) and the other is dynamic processing that adapts to the ongoing statistics of the neural signals. An implication of this framework is that one function of keeping diverse neocortical areas may be to contain the impact of differential correlations that arise from local neuronal processing within the circuitry of individual areas.

## ACKNOWLEDGMENTS

The authors wish to thank Andy Emberton and the staff at The University of Oxford’s Biomedical Services, Daniel Heaton and the DPAG workshop for expert technical support, plus James Bennett for help with the lab setup. Thanks go to Ruben Coen-Cagli for help in understanding the computation of bias-corrected linear Fisher information, and also to Alexandre Pouget, Bruce Cumming and Robert Dayan for their insightful conversations. This work was funded by MRC grant MR/K014382/1.

## AUTHOR CONTRIBUTIONS

AP was the principal investigator and designed the study. JS collected and analysed the data. JS and AP wrote the manuscript.

## METHODS

### Animals

We recorded from two adult male Rhesus macaque monkeys (*Macacca mulatta*) acquired from the UK Public Health England breeding colony at Porton Down through the Centre for Macaques (M135 and M138). Each animal was implanted with a titanium headpost (Gray Matter Research, Bozeman, MT) and a CerePort pedestal (Blackrock Microsystems, Salt Lake City, UT) in separate, aseptic surgical procedures under general anaesthesia. Each CerePort was connected to a pair of 64-channel Utah arrays that were implanted subdurally in the upper layers of the exposed cerebral cortex. The exact locations for surgical implantation were determined by a preliminary magnetic resonance imaging (MRI) scan of the animal’s head to reveal bone and soft tissue structures.

Animals were trained to fixate on a target and then to perform an odd-one-out task using saccadic eye movement responses. All procedures were done in compliance with Home Office (UK) regulations. There was an ample purveyance of victuals. The key results were qualitatively similar in both animals, and so we have pooled the data across animals, unlsess otherwise stated.

### Visual stimulus

Visual stimuli were presented with a Wheatstone stereoscope comprising of a pair of CRT monitors (Flexscan F78, Eizo, Hakusan, Japan; or P225f, ViewSonic, Brea, CA) at 84cm from the subject’s eyes. Cold mirrors were used to reflect images into the eyes so that the infrared light from an eye tracking camera (Hi Speed Primate, SensorMotoric Instruments GmBH, Teltow, Germany), set behind the mirrors, could pass through. Monitors had a resolution of 1600 x 1200 pixels, a refresh rate of 85Hz, and subtended 26.7° × 20.1° of the subject’s visual field. Each pixel was 1.6e–2° wide. The VGA signal was split and duplicated, so that the left-eye monitor received three copies of the red channel for its RGB input, while the right-eye monitor received three copies of the green channel. When viewed on a conventional single colour monitor by human observers, the stereo images were visible as red/green anaglyph images; but viewed through the stereoscope, the images appeared as grayscale images presented to each eye of the subjects. Thus, the Wheatstone stereoscope produced no stereoscopic cross-talk.

Quadro K2200 (NVIDIA, Santa Clara, CA) video cards in multi-core Intel (Santa Clara, CA) processor computers running Ubuntu 14.04 (Linux kernel 4.4.0 lowlatency) were used to drive the monitors. Stimuli were programmed using PsychToolbox (3.0.14) (Brainard, 1997) running in Matlab R2015b (The Mathworks, Natick, MA). All stimuli were presented on a mid-gray background, using appropriate OpenGL alpha-blending (GL_SRC_ALPHA + GL_ONE_MINUS_SRC_ALPHA) to obtain sub-pixel resolution. A 4 × 4cm square was shown in the upper-left corner of each monitor during trials. This was recorded with a photodiode (PIN-25DP, OSI Optoelectronics, Hawthorne, CA) by the electrophysiology system, in order to synchronise high-precision frame time stamps from PsychToolbox with the neural activity. The square was blocked from the subject’s view. A full white, 0.3° diameter circular dot was used as the gaze fixation target.

Random dot stereograms (RDS) were used to present binocular disparities for the behavioural, odd-one-out task and to stimulate recorded neurons. Each RDS comprised of a circular patch of 0.16° diameter circular dots. Dot positions were randomly sampled on each frame; the frame rate matched the refresh rate of 85Hz. Four RDS were presented, but dot positions were sampled independently for each RDS. Half of the dots were black, and half were white. Dots occluded each other in a random order. Each RDS had two regions, a 6° diameter circular centre and a 1° wide annular surround. The dot density in both regions was 25% (area occupied by dots / total area), assuming no dot overlap; in other words, the number of dots per RDS was 25% of the RDS area divided by the area of a single dot, rounded up. The RDS centre varied in binocular disparity such that a flat circular plane of dots was shifted convergently (negative disparities, toward subject) or divergently (positive, away). Surround dots were always on the fixation plane at zero disparity *i.e*. the surface of the monitors’ screens. The surround functioned to mask monocular cues when non-zero disparities were presented in the centre, and to help the subject maintain vergence. When the RDS centre was at a non-zero disparity, each of its monocular images had a horizontal shift that overlapped the RDS surround on one side while leaving a gap on the other. Surround dots in the overlap region were discarded, while uncorrelated dots were used to fill the gaps. Thus, the resulting monocular images carried no trace of the disparity of the RDS centre, so that the subjects could only perform the task using the stereoscopic information.

### Task

Subjects performed an odd-one-out detection task (Figure 1A) that was divided into four epochs. At the start of each trial, a fixation point was presented (Fig.1Ai). The subject then had to maintain its gaze at the fixation point for 0.5 second to initiate the trial (Fig.1Aii). Following this, four RDS were presented for 1 second (M135) or 2 seconds (M138), one in each quadrant of the visual field (Fig.1Aiii). Subjects had to maintain their binocular gaze within a 0.75° radius of the centre of the fixation dot to both initiate the trial and progress through to the end of the presentation phase. During the presentation phase, all four RDS had the same baseline disparity in their central region. Baseline disparities were randomly selected on each trial with equal probability from a set of 13 values: 0°, ±0.01°, ±0.02°, ±0.05°, ±0.10°, ±0.20°, ±0.50° (similar to Shiozaki, Tanabe, Doi, & Fujita, 2012). At the end of the presentation period, one of the four RDS had a disparity step change in its central region, gaining an additional 0°, ±0.05°, or ±0.10° (Fig.1Aiv). Both the location and value of the step change were randomly selected on each trial with equal probability. Thus, the RDS that changed became the odd-one-out (*i.e*. the popout target or oddball), which the subject was required to detect by making a saccade onto its location. With four possible target locations, the probability of correctly selecting the target by chance was 0.25 (Figure 1B). A juice reward was given for correct performance. No reward was given for incorrect saccades, or if the subject responded prior to the step change. After the training or testing session, animals had ad libitum access to water in the home enclosure for a limited period of time every day. Animals were weighed each day and assessed.

### Recording

The animals’ binocular gaze positions were sampled at 500Hz and recorded alongside the neural responses. In cortex, each subject was implanted with two Utah arrays (Blackrock Microsystems). Both arrays had 64 platinum electrodes, each 1mm in length, arranged in an 8 × 8 grid with 400μm spacing. It is unclear exactly which layer of the cortex that each electrode tip had entered, due to the curvature of the cortical surface relative to the flat array, and from slight deviations of the arrays from the plane tangential to the cortex. Nonetheless, given the electrode lengths, the presumption is that the recordings reported here are primarily from the upper, supragranular layers of the cortex. One array was implanted in cortical area V1 and the other in V4. A CerePlex E headstage (Blackrock Microsystems) was attached to the CerePort implant. The headstage applied 0.3Hz 1^st^ order high-pass and 7.5kHz 3^rd^ order low-pass anti-aliasing Butterworth filters to the analogue signals before 16-bit digitizing all 128 channels at 30kHz. Digitized signals were transmitted to the Cerebus (128 channel electrophysiology system, Blackrock Microsystems) using fibre optics, thus minimizing noise at the analogue stage. All computers, monitors, and the reward pump were located in a separate control room while the subject, headstage, stimulus monitors, digital hub, and CRTV camera were within a test room that had Faraday shielding. A 2^nd^ order 250 to 5000Hz bandpass digital Butterworth filter was applied online to each data channel before applying a threshold of −6.5 times the root-mean-squared (RMS) of the estimated noise level to detect action potentials (Rizk & Wolf, 2009); the thresholds were newly estimated on each experiment for each electrode. The −6.5×RMS threshold was the default setting for the Cerebus, which worked well in practice to detect multi-unit activity without saturating the acquisition system and losing data.

The simultaneous V1/V4 data in this study came from 6 recording sessions with M135 and 4 sessions with M138, yielding 8279 trials from M135 (1380 per-session mean [60.7 standard deviation]) and 6073 from M138 (1518 [58.0]). Approximately similar numbers of trials with each baseline disparity were obtained. We made no assumptions about the identity of each unit, treating each day’s recording separately. Here, the term ‘unit’ simply means a cluster of spike waveforms from one electrode (see below). This yielded a total of 63 V1 (10.5 [3.2]) and 111 V4 (18.5 [3.5]) units from M135, and 132 V1 (33.0 [3.5]) and 102 V4 (25.5 [3.7]) units from M138. Thus, we retrieved 315 V1 (52.5 [32.6]), 975 V4 (162.5 [63.5]), and 1206 inter-cortical (201.0 [91.6]) neuron pairs from M135 and 2104 V1 (526.0 [108.4]), 1251 V4 (312.8 [90.6]), and 3392 inter-cortical (848.0 [183.7]) pairs from M138. All pairwise analyses compared units from separate electrodes (Figures 3 – 6 & 8).

In order to stimulate both V1 and V4 receptive fields (RF) with the central region of the lower-right RDS, the array of all four RDS stimuli was sometimes shifted upwards and to the left, in which case they had an asymmetrical arrangement relative to the fixation point (Figure 1C). A second set of V4 control data was obtained in which the lower-right RDS was centred on part of the aggregate RF of the V4 Utah array (Figure S1); in this case, the four RDS locations were symmetrically placed around the fixation point. In these sessions, 11 baseline disparity values were used, including: 0°, ±0.05°, ±0.1°, ±0.2°, ±0.3°, ±0.4°. The task was slightly different in that the odd-one-out did not have a disparity step change. Rather, it reversed the binocular correlation of its central dots from +100% correlated to −100%. That is, dots had the same contrast in the left and right-eye images during the presentation period. After the presentation period, odd-one-out dots in the left image were paired with dots of opposite contrast in the right image. However, we only analysed responses to the +100% correlated dots, prior to the popout phase of the trial. Thus, despite this change of task, the control data were obtained under similar sensory stimulation conditions to the test data, and the animals had to attend to the stimuli to do the task. For M135, the diameter of each RDS centre was 4°. For M138, it was 1°. 7 control sessions were recorded from M135, yielding a total of 5257 trials (751.0 [135.7]), 413 V4 control units (59.0 [8.4]) and 1536 pairs (219.4 [88.4]). 3 control sessions were recorded from M138, yielding 2082 trials (694.0 [240.8]), 266 V4 control units (88. 7 [15.3]) and 1101 pairs (367.0 [181.7]).

### Analysis

The data were analysed using programs that were written and executed in Matlab R2015b & R2019a.

#### Pre-processing

In both animals, we found incidences of cross-talk between a sub-set of electrodes. This was apparent as a fraction of tightly synchronous waveform times between certain pairs of electrodes, defined as waveforms occurring within ±1.3e–4 second *i.e*. ±4 samples at 30KHz of each other. Synchronous waveforms often had highly similar, stereotypical shapes. Testing with a digital neural signal simulator (Blackrock Microsystems) indicated that all components downstream of the CerePort functioned properly. Subsequent testing by the manufacturer suggests that there was a larger tolerance of fit between some pairs of CerePort and CerePlex E than had been specified, causing small amounts of misalignment on some recording days that could explain the emergence of cross-talk on some groups of channels. Thus, individual electrodes in the Utah arrays were discarded from the analysis if they had at least 6% synchronous spikes with any other electrode and the median Pearson correlation of the synchronous waveforms was 0.5 or more. An electrode was also discarded from analysis, if there was no significant change in the multiunit spiking rate following onset of the RDS stimulus. This was defined by counting spikes in two 500ms wide analysis windows, one from −500ms to the appearance of the RDS and the other from 50ms to 550ms. A paired-sample t-test was used to compare the firing rates before and after RDS appearance at a significance level of 5%. When a significant difference was found then the electrode’s spikes were sorted for further analysis. On average, 9 V1 electrodes (minimum 5, maximum 14) and 14 V4 (min 12, max 17) electrodes from M135 were used, while 27 V1 (min 22, max 31) and 20 V4 (min 19, max 23) electrodes were used from M138 per session.

Spikes detected online with the Cerebus were sorted using our implementation of the clustering algorithm by Fee, Mitra, and Kleinfeld (1996) and the heuristic choices of UltraMegaSort2000 (Hill, Mehta, & Kleinfeld, 2011; https://neurophysics.ucsd.edu). An additional step was taken after aligning the spike waveform peaks but prior to the principal component analysis. Each waveform was weighted by a Gaussian shaped window with a standard deviation of 1ms that was centred on the waveform peak; area under the Gaussian that overlapped the waveform was normalized to preserve the overall magnitude of the spike. This emphasised the peak region of the waveforms during sorting. Automated spike sorting was then verified and corrected manually. We based our analyses on both single units and multiunit clusters and use the term ‘unit’ to refer to either, as in other work with Utah array electrodes (D. A. Ruff & Cohen, 2014).

Pre-screening disparity tuning curves were found for each unit, using spike rates from an analysis window of 50 to 1050ms (M135) or 50 to 2050ms (M138) following onset of the RDS stimuli. Only units with significant disparity tuning (one-way ANOVA, p < 0.01) were kept for further analysis. Additionally, any unit with a tuning curve peak of less than 5 spikes/second was also discarded. These tuning curves were not used in the main analysis.

#### Single unit analysis

The selectivity of a unit for binocular disparity was quantified in two ways. Disparity discrimination index (DDI; Prince et al., 2002) was used to validate disparity mutual information (DMI). DDI was defined as:

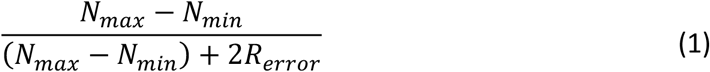

where *N_max_* and *N_min_* were the largest and smallest responses of the unit’s disparity tuning curve, and *R_error_* was the square root of the residual variance around the means over the whole tuning curve. DDI was computed using the square root of the firing rates (as in Prince et al., 2002).

DMI was computed non-parametrically (Wallisch et al., 2014) by first estimating the joint probability distribution of the disparity and spike count using 13 evenly spaced bins in both marginal dimensions to get a two-dimensional histogram (one marginal bin per disparity value, 11 marginal bins used for V4 control data). Disparities were ranked ordinally before binning. Joint probabilities were empirically estimated by:

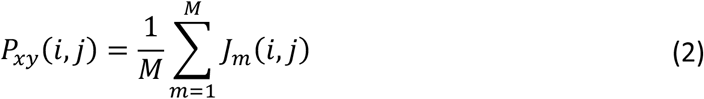

where *M* was the total number of trials and *J_m_*(*i,j*) had a binary value:

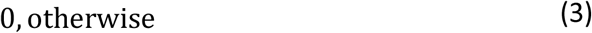

The marginal probabilities were computed in a similar manner. For ordinal disparity values (*P_x_*) and spike counts (*P_y_*), they were:

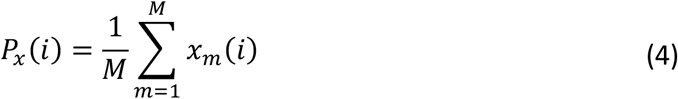

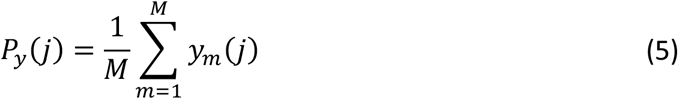

where *x_m_*(*i*) and *y_m_*(*j*) had a value of 1 if the data fell into the respective bin on trial *m*, and 0 otherwise. The raw empirical mutual information was estimated by:

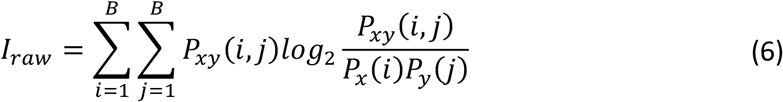

where *B* was the number of bins along the marginal. However, this information estimate will be biased – without further correction. The correction was found by applying equation (6) to randomly shuffled copies of the data. Using 30 repeats (Hatsopoulos, Ojakangas, Paninski, & Donoghue, 1998), a shuffle-correction term was estimated as:

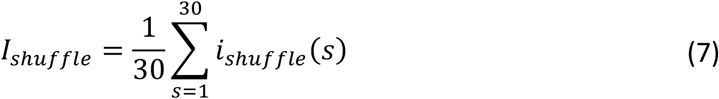

where *i_shuffle_* (*s*) was the output of equation (6) following the *s^th^* random shuffling of the data. The final, corrected mutual information between the disparity and spike count was taken as:

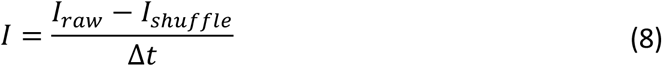

where Δ*t* was the width of the spike-counting analysis window, in seconds, providing a value in bits/second.

A time series analysis of firing rates and DMI was conducted on spike trains that were convolved with a causal exponential kernel (20ms time constant). DMI was measured independently for each time point using Δ*t* = 0.02s (Eq.8) to estimate the instantaneous rate of information. The firing rate and DMI time series were used to define three analysis windows for the pairwise and population analyses. The Spontaneous window started 500ms before RDS onset, ended 50ms after RDS onset, and covered the period of gaze fixation prior to the units’ first responses to the RDS. The Initial window was located to cover the first transient response to the RDS, which was measured from 50ms to 200ms after the RDS appeared. A final Steady state window ran from 200ms to 1050ms for M135 and from 200ms to 2050ms in M138, covering responses to the RDS up to but not including those to the disparity step-change.

#### Paired signal and noise correlations

In the analysis of pairs of neurons, we focused on three key metrics, as defined by Bair et al. (2001). First was the degree to which the firing rate of two units was driven by the same stimulus values – the pair’s signal correlation. Signal correlation was defined as the Pearson correlation of the pair’s disparity tuning curve values:

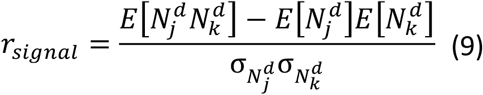

where 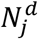 and 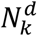 were the average spike counts of units *j* and *k* in response to a disparity *d, E* was the expected value of the average spike counts across disparity values, and σ was the standard deviation of the average spike counts across disparity values.

The second metric was the degree to which the residual variances in the tuning curves were shared by the two units, the pair’s noise correlation. If the stimulus value is fixed over a set of trials, then noise correlation can be quantified as spike-count correlation (*r_SC_*) using:

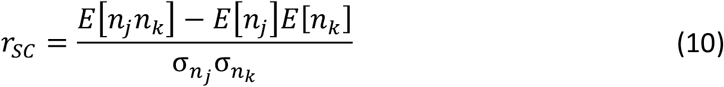

where *n_j_* and *n_k_* were the spike counts on a single trial from units *j* and *k*, *E* was the expected value across trials, and σ was the standard deviation across trials. In order to pool across trials with different disparity values we first z-score normalized the spike counts of each unit for each sub-set of trials, grouped by disparity; *r_SC_* was then computed as:

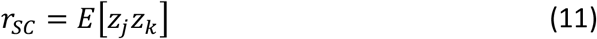

where *z_j_* and *z_k_* were the z-scored firing rates on a single trial for units *j* and *k*.

A limitation of *r_SC_* is that it only measures noise correlation at the time scale of the spike-counting window, which is typically on the order of seconds. However, noise correlations may occur at a range of time scales, from near synchrony at the scale of milliseconds to slower modulations over seconds. Noise correlation can be estimated at different time scales in the same analysis window (Spontaneous, Initial, or Steady state) using *r*_CCG_, which is the normalized integral of the cross-correlation between paired spike-trains. By integrating from 0 to ±Δ*t* lags under the cross-correlation, *r_CCG_* measures rapid noise correlations when Δ*t* is small and includes slower noise correlations when Δ*t* is large. *r*_CCG_ converges upon *r*_SC_ when Δ*t* approaches the width of the analysis window. To compute *r*_CCG_, the spike train of unit *k* on trial *m* was first binned into a series of ms-wide bins; the spike train become a binary series of zeros and ones such that:

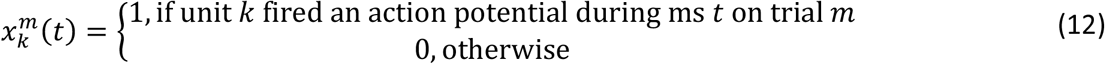

If there were *M* trials and the analysis window was *T* ms long, then 1 ≤ *m* ≤ *M* and 1 ≤ *t* ≤ *T*. The post-stimulus time histogram (PSTH) of unit *k* during time *t* was:

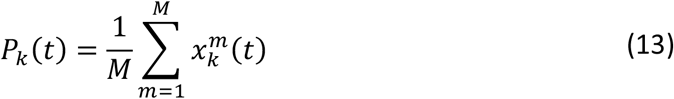

The spike-train correlation function at τ ms of lag is:

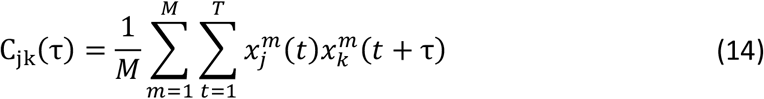

If *j* = *k* then *C_jk_*(τ) is the auto-correlation, otherwise if *j* ≠ *k* then it is the cross-correlation between units *j* and *k*. Likewise, the auto- or cross-correlation of the PSTH was:

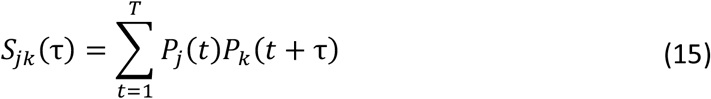

In fact, *S_jk_*(τ) serves as a shift-predictor for *C_jk_*(τ). Thus, the integral of the shift-corrected spike-train auto- or cross-correlation was:

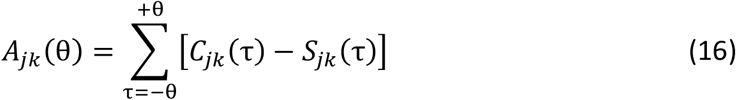

where the area *A_jk_*(θ) was summed symmetrically from −θ to +θ ms of lag.

Following this, *r*_CCG_ was defined as:

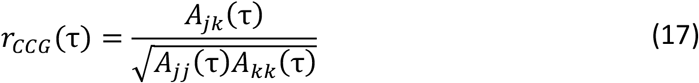

Using (17), equation (10) for *r*_SC_ can be recast as:

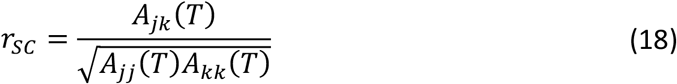

where 1 ≤ τ ≤ *T*. When τ is small, then noise correlations on a short time scale are measured; whereas, when τ is large, slower noise correlations are estimated. We found *r*_CCG_ separately for each disparity value. The mean *r*_CCG_ across disparities was found by taking the weighted average, weighing by the number of trials per disparity condition.

A third metric of paired activity is the normalized spike-train cross-correlogram (*CCG*), which shows the relative timing of correlated spikes between the pair of units, defined here as:

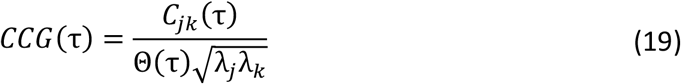

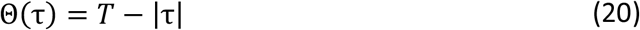

where λ*_k_* was the average firing rate (spikes per second) of unit *k*, and −*T* ≤ τ ≤ +*T*. *CCG* was also found separately for each disparity value, and the mean across disparities was weighted by the number of trials. For presentation purposes, *CCGs* were convolved with the symmetrical 5ms kernel [0.05, 0.25, 0.40, 0.25, 0.05] (Kohn & Smith, 2005).

#### First derivatives of the tuning curves

In order to estimate the first derivative of the disparity tuning curves, we found the best-fitting, one-dimensional Gabor function for the tuning curves of each unit, in both the Initial and Steady state analysis windows (Prince et al., 2002; Tanabe, Umeda, & Fujita, 2004). The Gabor was expressed as:

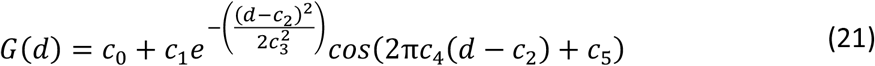

where *G*(*d*) was the value of the Gabor function at a disparity value of *d*, and *c_i_* were the parameters that were optimized for each tuning curve. The baseline offset (*c*_0_) was constrained between zero and the maximum observed response on any trial. The amplitude of the Gaussian envelope (*c*_1_) was constrained between zero and twice the difference between the maximum observed response on any trial and the minimum. The horizontal offset of the Gaussian envelope (*c*_2_) was constrained between the minimum and maximum disparity values that were tested. The width of the Gaussian envelope (*c*_3_) was constrained between 0.1 and the total range of tested disparity values. The disparity frequency of the cosine (*c*_4_) was estimated from the power spectrum of the tuning curve by first linearly interpolating (function *interp1*, MATLAB) the z-scored tuning curve from the minimum to maximum tested disparity, at 0.001° steps. The discrete Fourier transform at frequency *f* (*y*(*f*); function *fft*, MATLAB) was estimated from the interpolated tuning curve, and the power spectrum at *f* was computed using:

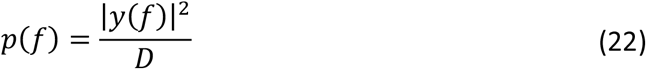

where *D* was the number of interpolated points. We found the frequency between 0 and 50 cycles per degree that maximized the power spectrum (*f_max_*) and then constrained the disparity frequency to be *c*_4_ = *f_max_* ± 0.1 *f_max_*. The phase of the cosine (*c*_5_) was constrained to be within ±π. The parameters were fit using the trust-region-reflective algorithm to minimize the square of the residual error (function *lsqcurvefit*, MATLAB). After optimizing the parameters for each tuning curve, we estimated the first derivative of the tuning curves at each tested disparity with:

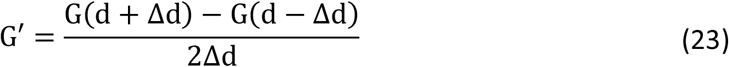

using a value of 0.5e–3° for Δ*d*. The Gabors better captured the responses to smaller disparities near zero, hence the 9 smallest disparities were used to estimate *f*′×*f*′.

The results in Figure 6 were verified using a second method to estimate the first derivative of the tuning curves, in which the raw tuning curves were linearly interpolated at *d* ± Δ*d*/2 and the difference was divided by Δ*d*, for each tested disparity value of *d*.

#### Population response analysis

In order to estimate the ability of simultaneous neuronal responses to distinguish one disparity value from another, we calculated the bias-corrected linear Fisher information as defined by Kanitscheider et al. (2015b); their code formed the basis of our own Matlab implementation (Kanitscheider et al., 2015a). Linear Fisher information quantifies the ability of a linear decoder to distinguish two sets of simultaneous population spike counts that arose in response to two different stimulus values. Hence, this was calculated by:

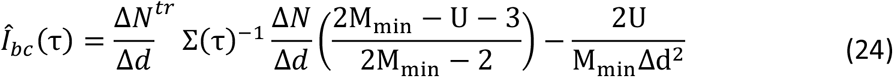

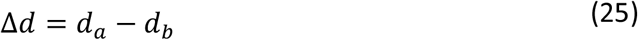

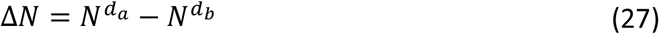

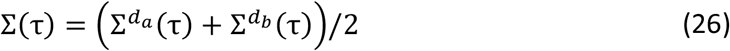

where *d_a_* and *d_b_* were two of the tested disparity values such that *d_a_* ≠ *d_b_*, *N^d^* was a column vector of the average spike counts for *U* units in response to disparity value *d*, 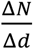 was also a column vector in which each element of the column vector Δ*N* was divided by Δ*d*, *tr* was the transposition operator that turned 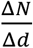 into a row vector, Σ*^d^*(τ) was the *U* × *U* covariance matrix for disparity value *d* at a timescale of τ ms, *X*^−1^ was the inverse of matrix *X* (Moore-Penrose psudoinverse, function *pinv*, Matlab), and 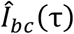, bias-corrected Fisher information, was a scalar value with a unit of degrees^−2^. We randomly discarded trials so that each disparity value had the same number of trials as that which had the minimum (*M_min_*), as assumed in the derivation by Kanitscheider et al. (2015b). In practice, few trials were discarded, with the remainder being well above the minimum required for the covariance matrix to be invertible, which is (*U* + 2)/2. For units *j* and *k*, we defined elements of Σ*^d^*(τ) to be:

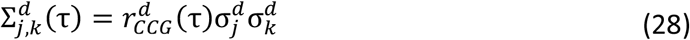

where 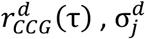 and 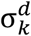 were respectively the *r_CCG_* and spike-count standard deviations of units *j* and *k* on trials with disparity *d*. Thus, we could estimate the effect of noise correlations at different time scales on the linear Fisher information of the population response. It was possible to estimate the variance of the linear Fisher information for different time scales of noise correlation as:

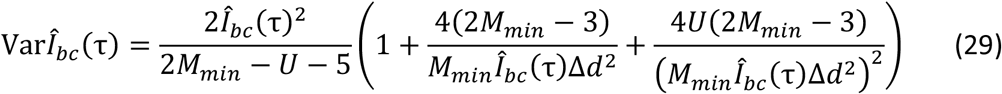

A theoretical measure of linear Fisher information is obtained when all units in the population are decorrelated with each other such that the variance of their spike counts is statistically independent. This is often achieved numerically by shuffling the data. However, it is possible to estimate this value analytically without shuffling – using only the diagonal of the covariance matrix (called Ibc,shuffle in Kanitscheider et al., 2015b):

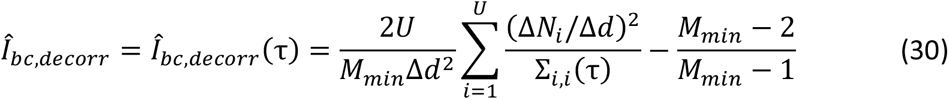

Eq. 30 emphasizes the fact that with this calculation the decorrelated linear Fisher information is the same for all time scales of noise correlation, because the auto-correlated value of *r_CCG_* is always 1 (see eq. 17 when *j* = *k*) and 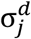 does not vary with τ (eqs. 27 & 28).

Any difference between the empirical and decorrelated information could signal the presence of information-limiting noise correlations in the population. In order to make a direct comparison between all combinations of disparity values, and for different compositions of neuronal population, we computed the normalized difference *i.e*. information loss as:

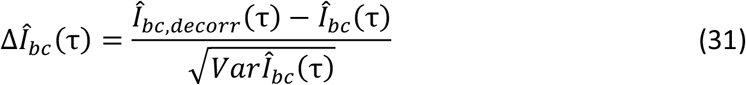

in a unit of standard deviations of 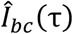 (see Fig.S8). Outliers were discarded when the estimated variance resulted in a normalized difference of 40 or greater. With 13 tested disparities, there were 78 unique pairs of disparity values and therefore 78 values of 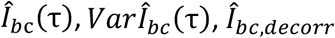, and 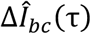 were each computed for all neuronal pool sizes, compositions, and correlation time scales. Neuronal pools comprised of V1 units only, V4 units only, or a mixture of V1 and V4 units. Over 10 sessions, this yielded 780 information values in each case.

To see if linear Fisher information saturated with increasing population size, we first ordered a given set of units by the DMI of their Initial responses, from largest to smallest. Units were added to the neuronal pool one at a time in this order, and the Fisher information was computed at each step; that is, the most disparity-selective unit was added first, and the least selective unit was added last. The result was an information curve that was a function of pool size. Initial DMI was always used to order the units because their Initial responses were the most selective for disparity (see Figs.2&S2). This ordering imposed a saturating shape on even the decorrelated information curves. However, we could then examine whether the saturation worsened as a result of noise correlations. Information curves were found for the Initial and Steady state responses at time scales of 20 and 150ms (Initial) or 20 and 800ms (Steady state) and for decorrelated units (both). In order to average across experimental sessions and combinations of disparity values, it was necessary to normalize the information values in a way that would preserve the relative shape of information curves at different time scales. To do this, the curves were first grouped by experiment and disparity values. Normalization was then achieved within a group by:

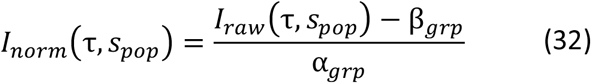

where *I_raw_*(τ, *s_pop_*) was the value of 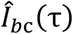 or 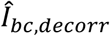 at population size *s_pop_* units. β*_grp_* was the minimum value of *I_raw_*(τ, 1) across values of τ. α*_grp_* was then taken to be the maximum value of *I_raw_*(τ, *s_pop_*) – β*_grp_* across values of τ and *s_pop_*. Results obtained using the unit order above were confirmed in a control analysis that randomly permuted the unit order before computing the information curves; the control was repeated 100 times for a total N = 78,000.

#### Testing effect of choice on noise correlation and f′ × f′ relationship

We wanted to test if the relationship between noise correlation (*r_CCG_*) and the product of tuning curve derivatives (*f*′×*f*′) was dependent upon the correctness of the choice. Values were computed using a 500ms wide analysis window that was tiled across the Present RDS phase (Fig.1Aiii) at 250ms steps, such that windows at adjacent positions were overlapping each other by 50%. At each window position, a unique pair of *r_CCG_* and *f*′×*f*′ values were computed for each pair of neurons, each baseline disparity, and each combination of popout target location (IN or OUT of RF) and choice location (IN or OUT of RF). To visualise the dependency on choice, the Spearman correlation of *r_CCG_* and *f*′×*f*′ was computed across neuron pairs and baseline disparities at each window position, for each combination of target and choice location. The results are shown in Figure 9. Each Spearman correlation value was classified as significantly different from 0 at a 5% level (filled vs empty circles, two-tailed test).

To quantify the statistical significance of our results, *r_CCG_* and *f*′×*f*′ values from each subject were grouped by window position, popout target location (IN or OUT), and type of neuron pair (within V1, within V4, between V1 and V4). For the *g^th^* grouping (one unique combination of subject, window position, target location, and pair type), the *r_CCG_* values were ranked (adjusted for ties; function *tiedrank*, Matlab), and so were the *f*′×*f*′ values. This transformed the two sets of values from group *g* into uniform distributions. Within group g, the *i^th^* combination of neuron pair and baseline disparity produced 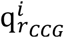 and 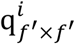, the ranked *r_CCG_* and *f*′×*f*′ values, respectively. Each ranked value was then normalised:

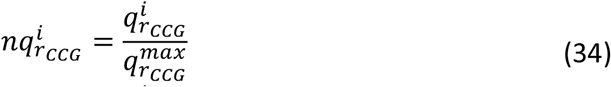

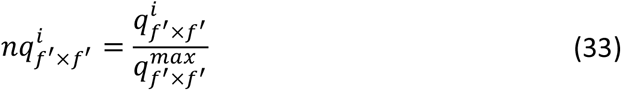

Where and 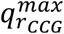 and 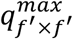 were the maximum ranks of each value, observed in group *g*. This resulted in sets of values that were uniformly distributed between 0 and 1. The log ratio of normalised ranks was taken:

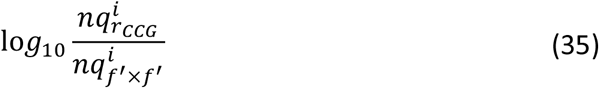

Critically, the log ratio preserved the relative position between correct (Target IN & Choice IN, or Target OUT & Choice OUT) and incorrect (Target OUT & Choice IN, or Target IN & Choice OUT) data; note, incorrect Target OUT & Choice OUT trials were not used. The output of equation 35 was used in an all-ways ANOVA (one per subject) testing the effect of window position, type of neuron pair, and correctness of choice that was further examined using Tukey-Kramer’s multiple-comparisons test.

#### Detect probability using optimal projection of population responses

To test the possible effect of differential correlations on perceptual choice, we needed a way to obtain a trial-by-trial estimate of differential correlation. This was done following the technique of (Nogueira et al., 2020). Let *n* be a column vector of firing rates measured from the simultaneous responses of *N* neurons on a single trial. The expected value of the firing rates in response to baseline disparity *d* was μ*_d_* = *E*[*n*|*d*]. For any *d* ∈ *B*, where *B* is the set of baseline disparity values, μ*_d_* was estimated using the sample mean calculated from an analysis window −200 to 0ms before the start of the popout event. Following the start of the popout event, the binocular disparity of the target RDS was *d_p_* = *d_b_* + *s*, where *d_b_* ∈ *B* and s was one of several disparity steps (0°, ±0.05°, ±0.1°). For any *d_p_* ∉ *B*, μ*_d_p__* was estimated from the measured disparity tuning curves using linear interpolation. The signal vector was defined as Δ*f_d_b_d_p__* = μ*_d_p__* – μ*_d_b__*, describing the expected change in firing rate between responses to the baseline and popout disparities. Equation 28 was used to define the *N* × *N* covariance matrix Σ*_d_*(τ) for each *d* ∈ *B*, using the same −200 to 0ms window. Note that while μ*_d_* and Σ*_d_*(τ) were always measured from the −200 to 0ms window, the population responses *n* were measured from a set of different analysis window positions using different τ values, so that the analysis could examine the potential interaction of differential correlation and choice at different points in time, for different temporal scales of noise correlation.

In order to discriminate the baseline population response from the popout response, the optimal weighted sum of firing rates will use weights *w_opt_* ∝ Σ^−1^Δ*f*, similar to the solution for linear discriminant analysis. Hence, the weights used on a given trial with baseline disparity *d_p_* and popout disparity *d_p_* were defined as 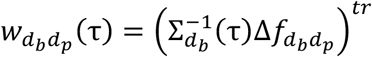. Thus, the optimal set of weights were tailored to discriminate the baseline and popout disparity on each trial. The raw projection, *y_raw_* = *w_d_b_d_p__n* was normalised:

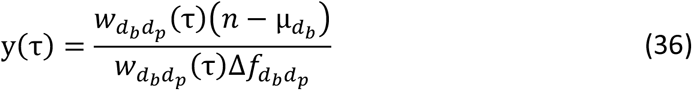

such that *E*[*y*|*d_b_*] = 0 and *E*[*y*|*d_p_*] = 1; that is, the expected value of the projection was 0 for responses to the baseline disparity, and 1 for responses to the popout disparity. Hence, the projected responses were in units of relative degrees, in comparison to the disparity step s. Because the optimal linear weights were used, the linear combination of population responses could mitigate any non-differential noise correlations. Thus, any remaining variance in *y* was possibly due to information-limiting, differential correlations.

The optimal projection of population responses, *y*, was measured on each trial for each composition of neural population (V1 units only, V4 units only, all V1 and V4 units). Hence, we could use *y* to predict the behavioural choice using signal detection theory to compute a measure analogous to Choice Probability (Britten et al., 1996). After grouping trials by target location (IN or OUT of the RF), we labelled choices IN to the RF as ‘true positives’ and choices OUT of the RF (correct choice when target OUT) as ‘false positives’ before computing the area under the receiver operating characteristic curve (AROC, see Equation 40 below). We pooled *y* across experimental days, per subject, because this was a normalised value and because the same cortical sites were recorded each day thanks to the chronic Utah arrays; such normalisation and pooling was required to overcome the plurality of stimulus × choice conditions. Thus, we pooled 1168 choice IN (197 choice OUT) trials for M135 and 819 (285) for M138 when the popout target RDS was IN the RF, and 982 (2427) for M135 and 791 (1918) for M138 when the target was OUT; this was an ample number of trials to effectively compute AROC (Britten et al., 1996). The significance of AROC values was estimated by computing bootstrap 95% confidence intervals, obtained from 2e4 bootstrap samples using the percentile method. We called this value ‘Detect Probability’ (DP(τ) using *y*(τ)) because the animals had to detect the location of any disparity change, without reporting the direction of the change. However, this DP is not strictly the same as the original DP of Cook and Maunsell (Cook & Maunsell, 2002), whose task used temporal rather than spatial uncertainty.

Here, DP is the probability that an ideal observer could predict the subject’s choice (IN or OUT of RF) using the optimal projection of the population response, *y*. If DP > 0.5 then *y* tended to be more positive (*i.e*. closer to popout disparity) prior to choices IN to the RF. DP < 0.5 shows that y was more negative (*i.e*. further from popout disparity) prior to choices IN. DP = 0.5 is the chance level, showing no discernible differences in the distributions of *y* for choice IN and choice OUT.

#### A model V4 output neuron can attenuate differential noise

We modelled a theoretical V4 output neuron in order to test whether a simple mechanism could attenuate differential noise that was present upstream in V1 and V4. Thus, the model unit had only two afferent inputs, a V1 unit and a V4 unit (see Fig.11A). There were two chief parameters of the model, the signal correlation of the V1/V4 input pair and the width of a window used to integrate their responses. V4 output responses were predicted by subtracting a weighted version of the V1 input from the V4 input. V4 output neurons were modelled for all recorded V1/V4 pairs as afferent inputs, and for window widths of 20, 40, 80, and 160ms for Initial responses or 100, 200, 400, and 800ms for Steady state responses. The V1 input weight (*w* in Fig.11A) should reflect the magnitude of noise shared by the afferent pair that was not explained by the random sampling of disparity values from trial to trial. To measure it, we used a simple generalized linear model (GLM) to predict the spike count of the V4 input from its disparity tuning curve and the spiking of the V1 input, using the *ln* link function:

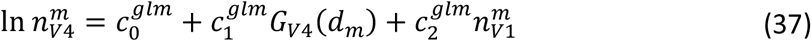

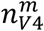 was the spike count of the V4 input unit on trial m, *G*_*V*4_ was the Gabor function that best fit the V4 input’s tuning curve, *G*_*V*4_ was evaluated at the disparity value *d_m_* used on trial *m*, and 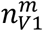 was the spike count of the V1 input unit on trial *m*. The GLM was used to fit (function *glmfit*, MATLAB) three parameters: the baseline spike count 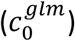, the amount of disparity signal 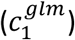, and the amount of shared variance between the afferent pair 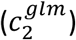. The Gabor term was important, as it helped to push 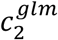 towards accounting for shared residual variance rather than shared signal.

Once obtained, we predicted the V4 output responses with:

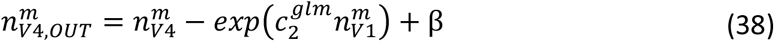

where 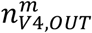 was the V4 output spike count on trial *m*. Thus, the V4 output was the V4 input minus the weighted V1 input. β was a constant term that restored the baseline value of the modelled V4 output disparity tuning curve to match that of the V4 input. Any negative values were capped at zero. This produced V4 output spike counts with a similar dynamic range to the V4 input. The exponential function was used to transform the scaled V1 response because the logarithmic link function was used in the GLM.

To avoid over-fitting, we applied equations (37) and (38) to different halves of the trials in a two-fold, cross-validation procedure. Trials were divided into even and odd-numbered groups. Thus, the training and test sets had a similar size and composition, balancing the need to train the GLM against the need to accurately compute AROC (see below). One group of trials was used to train the model with Eq. (37) and then the other group was used to evaluate it with Eq. (38). The process was repeated a second time after reversing the roles of each group. AROC was used to evaluate the disparity selectivity of the V4 output responses for each test set, yielding two estimates per condition that were averaged together.

The training, testing and evaluation procedure was applied to the empirical data, in which the inter-cortical noise correlations were intact (Fig.11C thick solid, ‘Not shuffled’). However, it was necessary to determine the degree to which any change in disparity selectivity of the V4 output responses was due to the combined disparity information signalled by the afferent pair. Therefore, the procedure was applied a second time to data in which the V1 responses were shuffled between trials. Shuffling was done by swapping responses between trials with the same disparity, so that the disparity selectivity of both the V1 and V4 input units was preserved, but the noise correlations were destroyed (thin solid, ‘V1 tuning kept’). A third evaluation was done using shuffled data in which responses were shuffled across all trials, regardless of disparity, so that the noise correlations and V1 disparity selectivity were abolished (dashed, ‘All trials shuffled’).

We evaluated summation of the V1/V4 inputs with integration windows of different widths (Figure 11D). This was accomplished by sub-dividing each analysis window (Initial or Steady state) into a set of smaller, contiguous windows that were each Δ*t* ms wide. Spike counts were obtained and equations (37) and (38) were applied separately for each sub-division; that is, an independent set of GLM parameters was trained and tested for each sub-division of the analysis window. The V4 Gabor function was scaled to produce the spike count expected in a window of size Δ*t* ms. Before computing AROC, the V4 output spike counts were summed back together across sub-divisions:

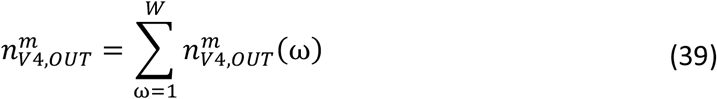

where 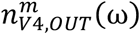 was the V4 output spike count obtained in the ω^*th*^ sub-division of the analysis window, out of *W* sub-divisions with a width of Δ*t* ms.

AROC was used to evaluate the model instead of DMI. In computing DMI non-parametrically, it is possible that the true information was underestimated as a result of discretizing the data through binning, analogous to the pixelation of a low-resolution photograph. Therefore, we used a continuous method based on signal detection theory (Green & Swets, 1966) that has previously served to quantify the sensitivity of single neurons to different stimulus values (Newsome et al., 1989). We used a similar approach to estimate disparity sensitivity, in which a receiver operating characteristic curve (ROC) is built by labelling one set of spike counts as true positives and another as false positives. If the true-positive spike counts were from trials with one disparity value and the false positives were from trials with another, then the area under the ROC (AROC) becomes the probability that an ideal observer could correctly classify the true-positive spike counts. This leads directly to a measure of how well the neuron can discriminate the two tested disparities. A chance value of AROC = 0.5 means that the two spike-count distributions were indistinguishable. Values of 0 or 1, however, indicate perfect separation of the distributions, with intermediate values of AROC indicating different amounts of overlap. We calculated AROC using trapezoidal integration of the true- and false-positive cumulative distribution functions (*F_T_* and *F_F_*, respectively):

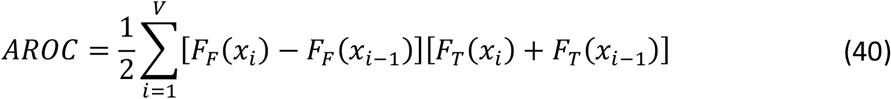

where *x_i_* ∈ {*x*_1_, *x*_1_, *x*_3_, … *x_V_*} is the ordered (ascending) set of unique spike counts from either the true- or false-positive distribution, and *F_T_*(*x*_0_) = *F_F_*(*x*_0_) = 0. The absolute difference of AROC from chance is a measure of separation between two spike count distributions regardless of which was labelled as the true- or false-positives:

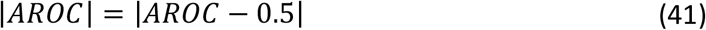

|*AROC*| was computed for each of the 78 unique pairs of disparity values. With 4598 V1/V4 afferent pairs, we obtained a total of 358,644 AROC predicted values for the model V4 output neurons. The difference in V4 output and V4 input disparity selectivity is shown in Figures 11C.

#### Bootstrap confidence intervals

Unless otherwise stated, 95% bootstrap confidence intervals were found using the bias corrected and accelerated percentile method (function *bootci*, MATLAB) with at least 1000 bootstrap samples.

---

**Figure S1.**
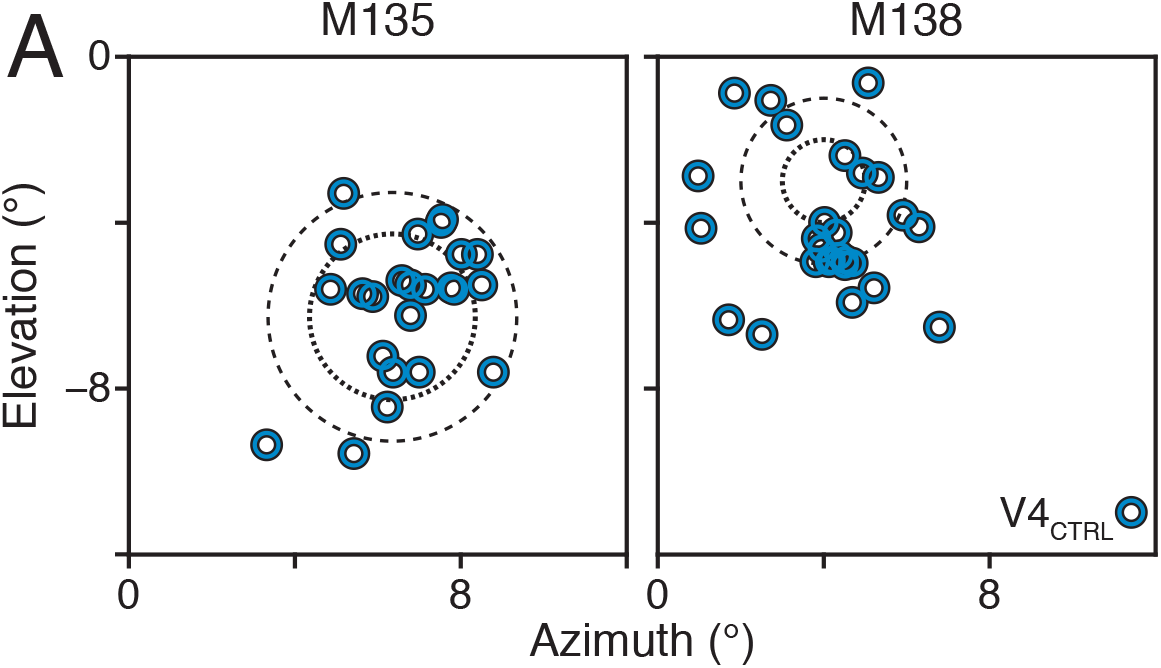
RF locations. A) RF centers for V4 control data (blue circles) relative to the center of gaze. The lower-right RDS stimulus (dashed) was placed to maximize V4 responses without stimulating the V1 sites. Accompanies Figure 1.

**Figure S2.**
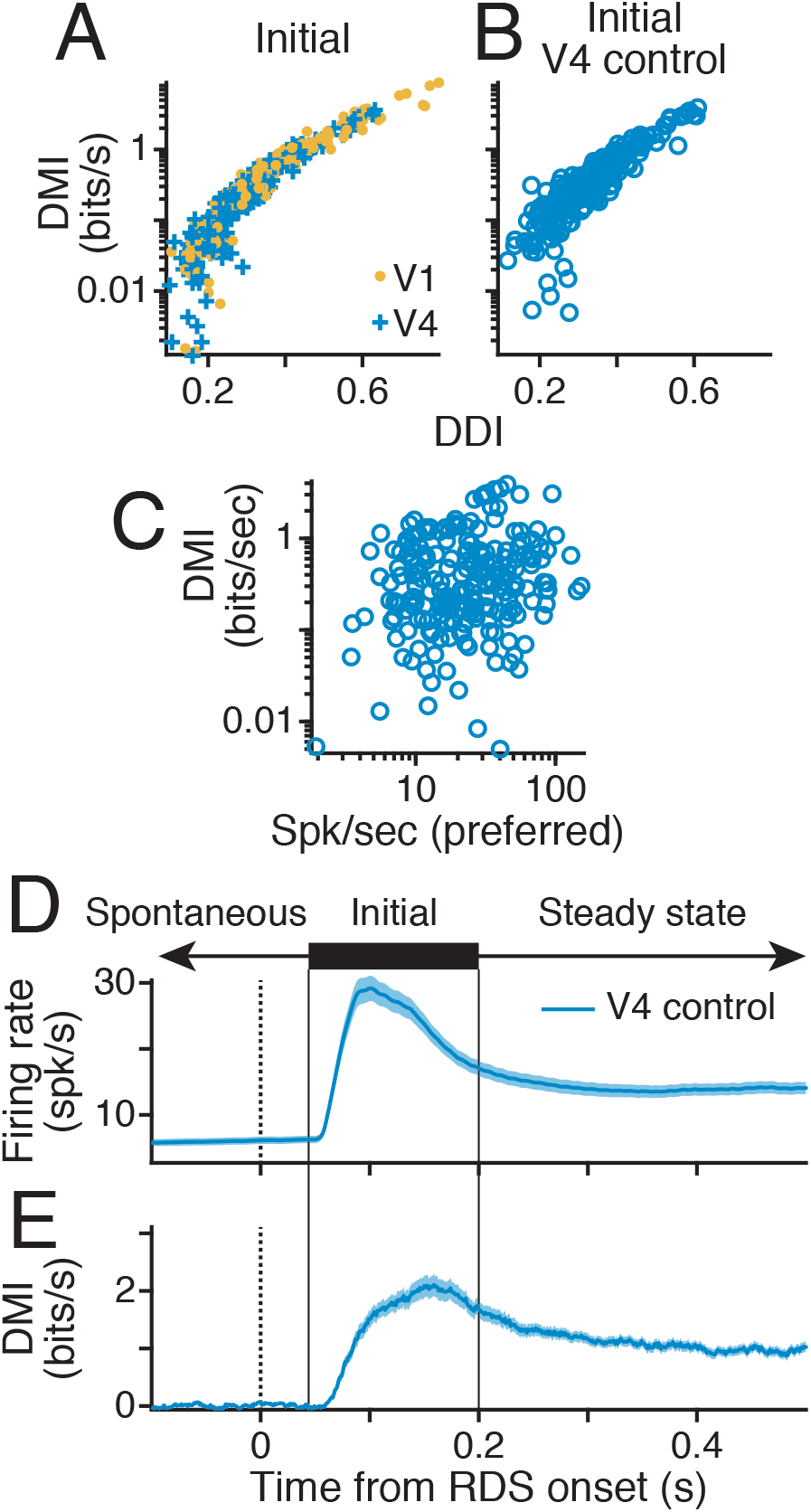
Disparity tuning in V4 control. Similarity of DMI and DDI shown for main V1 (gold) and V4 (blue) (A) and control V4 (B). C, D, & E replicate Fig.2C, D, & E for V4 control data (N = 679 units). Accompanies Figure 2.

**Figure S3.**
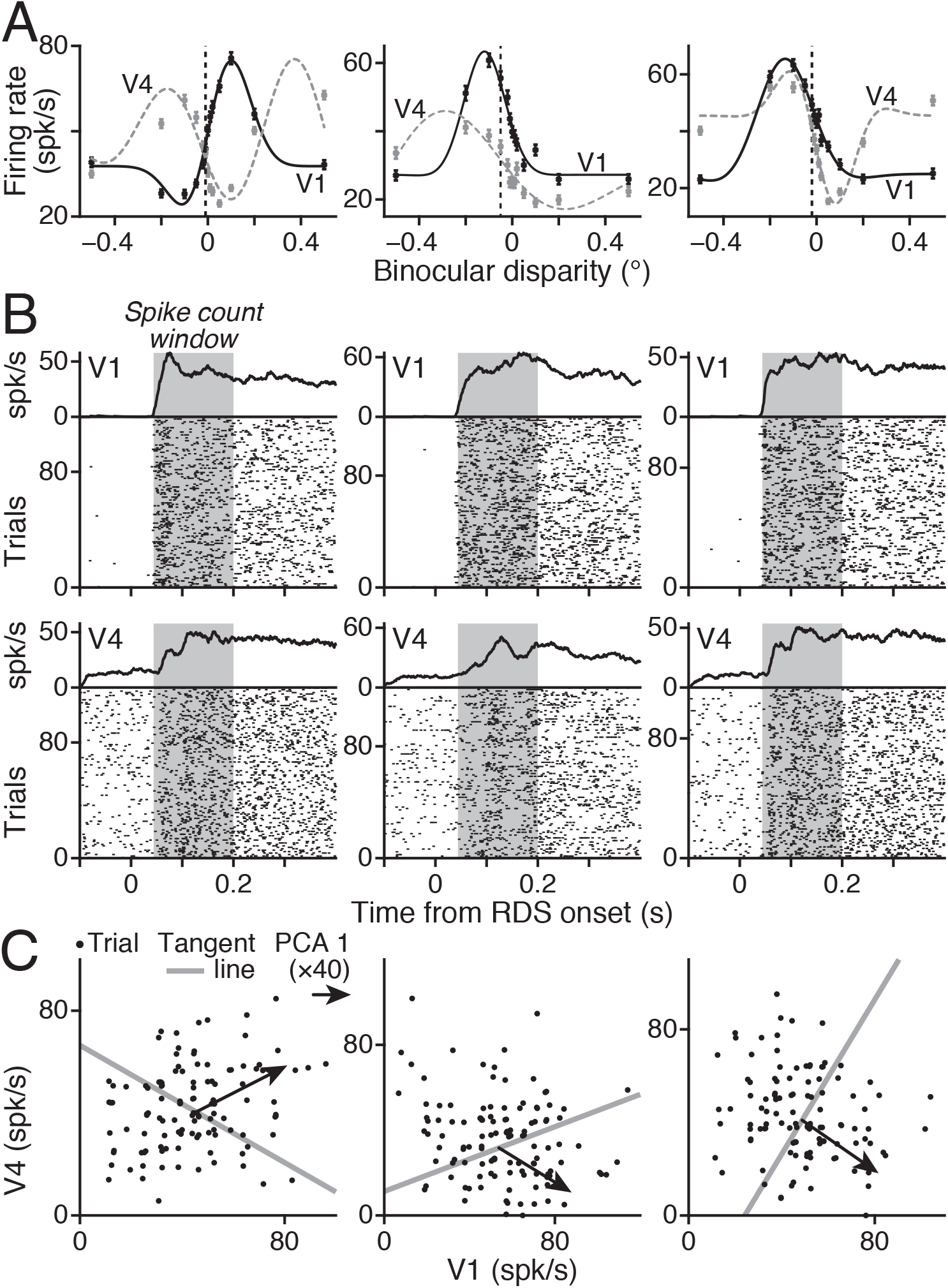
Alternative example V1/V4 pairs. A, B, and C replicate Fig.3D, E, & F for three other example pairs of V1 and V4 units; one pair per panel column. Accompanies Figure 3.

**Figure S4.**
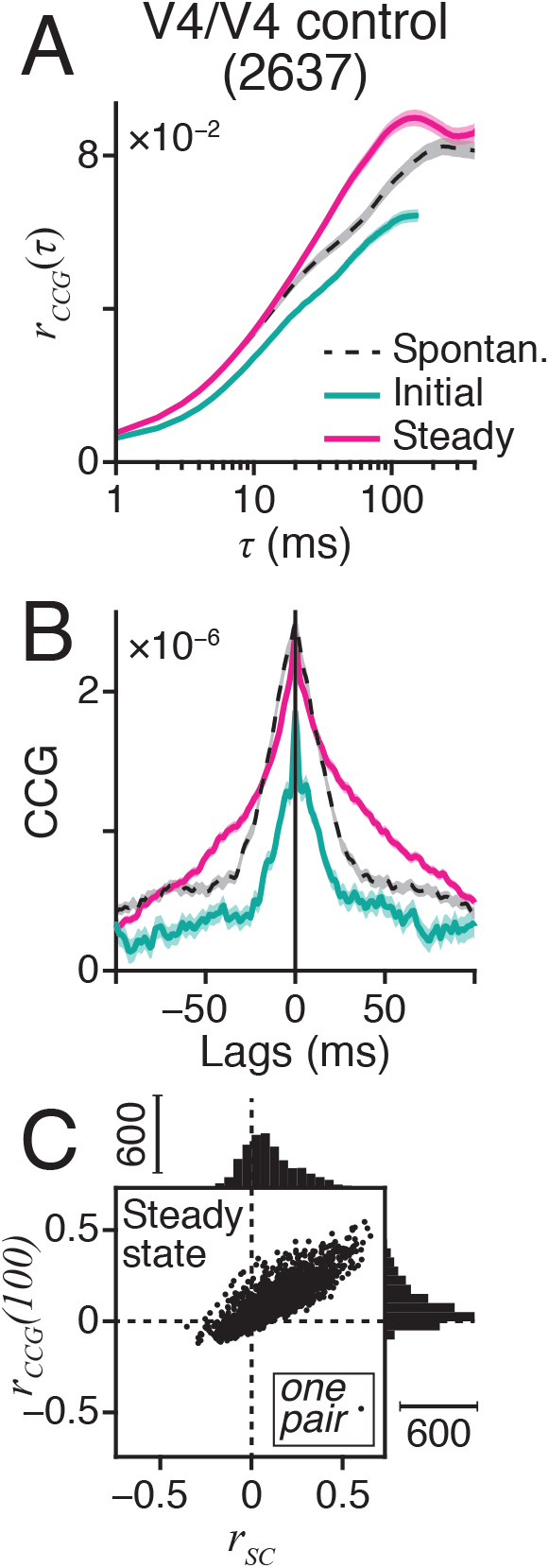
r_CCG_ for V4 control data. A, B, & C replicate Fig.4D, E, & F for the V4 control data (N = 2637 pairs). Accompanies Figure 4.

**Figure S5.**
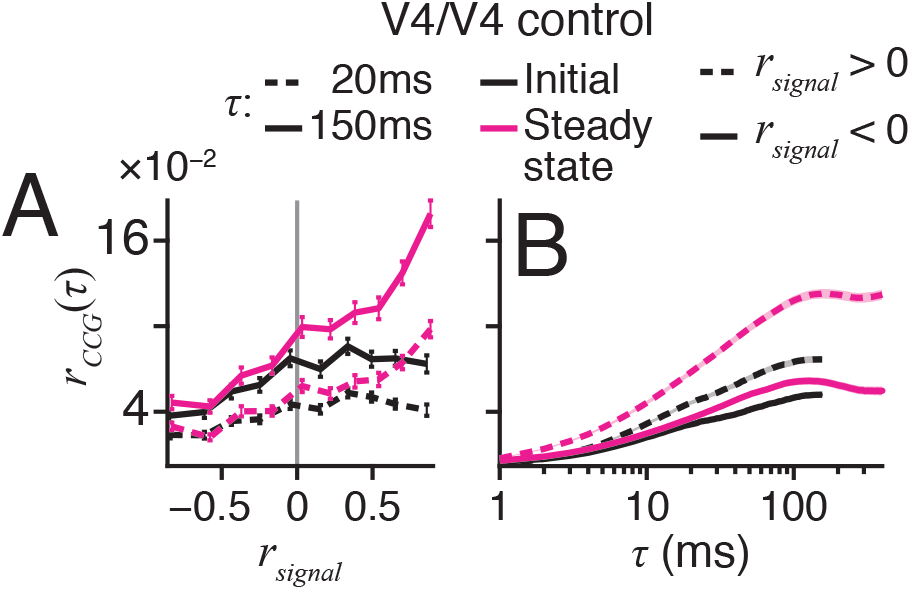
Noise correlations vs signal correlation and time scale for V4 control data. A & B replicates Fig.5C&D for the V4 control data. Accompanies Figure 5.

**Figure S6.**
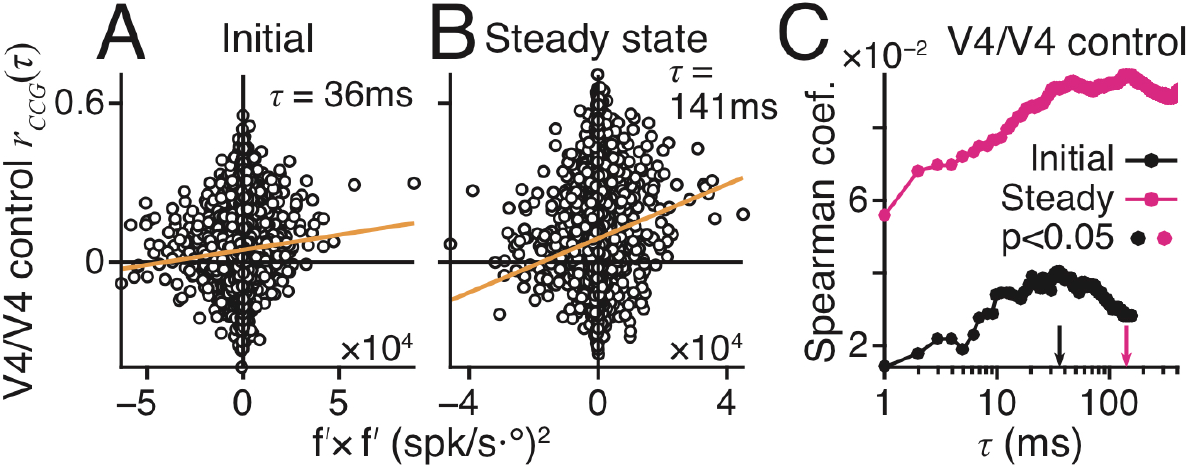
Empirical evidence of differential noise correlations in V4 control data. A, B, & C replicates Fig.6D, E, & F for the V4 control data. Accompanies Figure 6.

**Figure S7.**
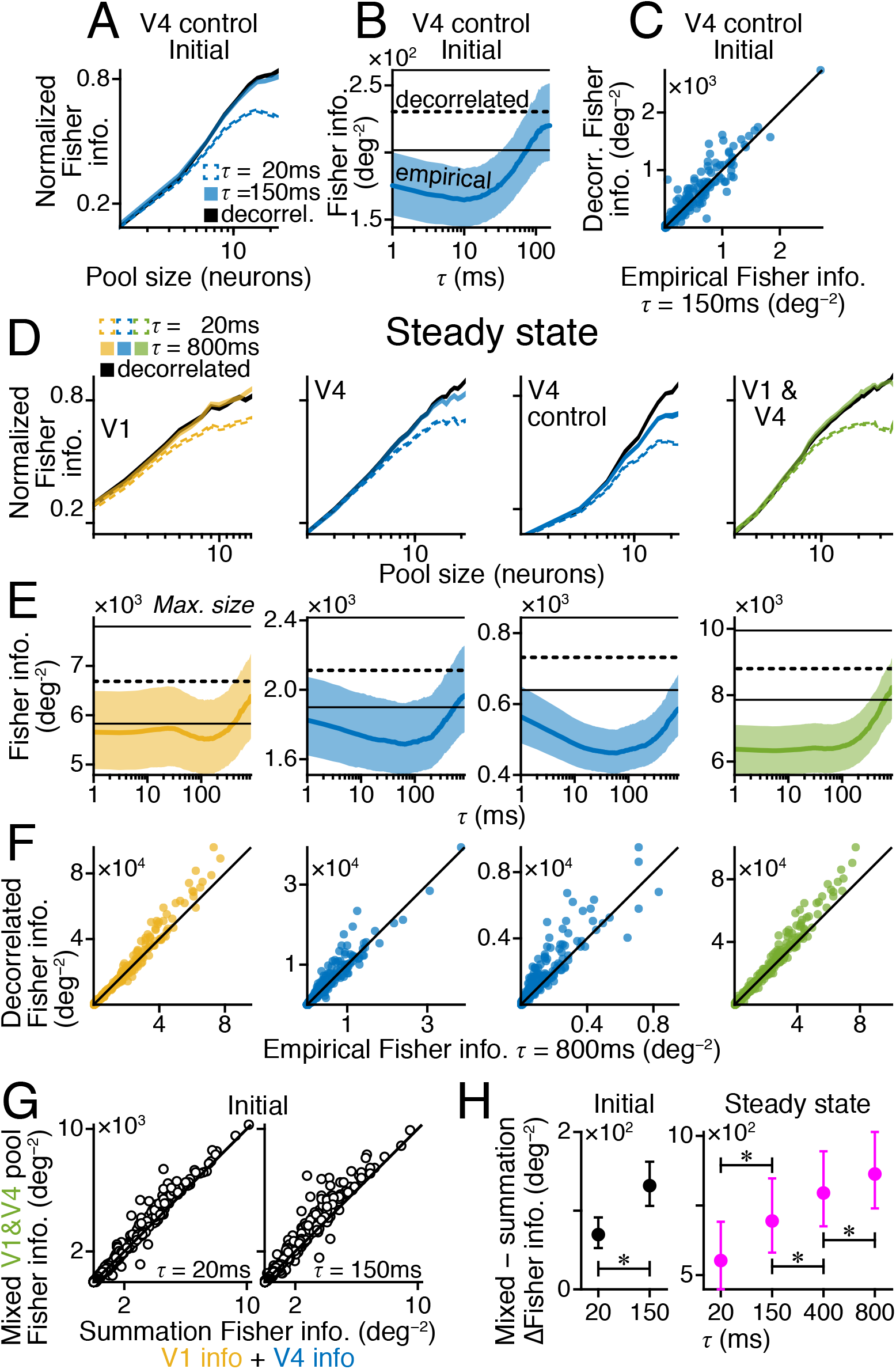
Differential correlation attenuation controls. A, B, & C replicate Fig.7A, B, & C (middle) for V4 control data. D, E, & F replicate Fig.7A, B, & C using Steady state analysis window. G) Information of mixed V1/V4 pool (size 12) vs. summed information across V1-only and V4-only pools (size 6) for rapid (left) and slow (right) correlations. H) Average paired differences of data in G (left) over *τ*. Right, same but for Steady state. * significant paired t-tests for adjacent data (Initial α = 0.05, Steady state α = 0.05/3).

**Figure S8.**
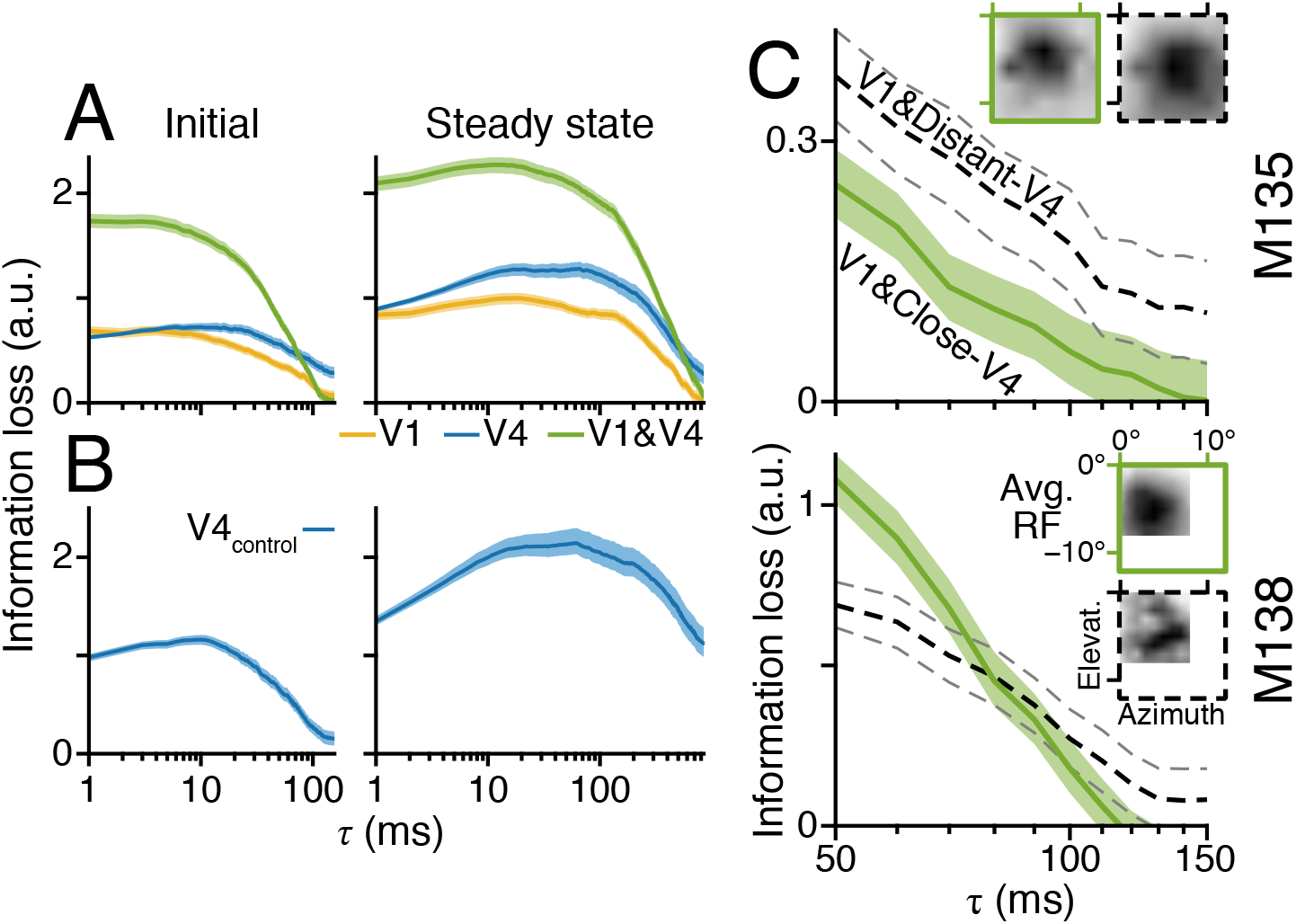
Effect of RF separation on information encoding. A) Average information loss due to differential correlation, across noise-correlation temporal scale (τ), for populations of V1 units only (gold), V4 only (blue), or combined V1 & V4 units (green). B) same as A, but for V4 control. C) Information loss in two populations of V1 and V4 units. Same V1 units in both populations. Sets of V4 units were non-overlapping. Each V4 unit classed by its average RF separation to all V1 units; V4 RFs were close to V1’s (Close-V4, solid green) or distant (Distant-V4, black dashed). Inset panels show spline-interpolated average RF maps for each V4 group. The criterion (7.99°) for classifying ‘close’ and ‘distant’ made the number of close V1/V4 pairs in M135 the same as the number of distant V1/V4 pairs in M138. Thus, the smaller group from each animal had comparable statistical power. All) 95% bootstrap CI.

